# The Development, Function, and Plasticity of the Immune Macroenvironment in Cancer

**DOI:** 10.1101/805473

**Authors:** Breanna M. Allen, Kamir J. Hiam, Cassandra E. Burnett, Anthony Venida, Rachel DeBarge, Yaron Carmi, Matthew H. Spitzer

**Author notes:** Co-First Author (Breanna M. Allen, Kamir J. Hiam). Correspondence (M.H.S.).

## Abstract

Harnessing immune defense mechanisms has revolutionized cancer therapy, but our understanding of the factors governing immune responses in cancer remains incomplete, limiting patient benefit. Here, we use mass cytometry to define the organism-wide immune landscape in response to tumor development across five tissues in eight tumor models. Systemic immunity was dramatically altered across mouse models and cancer patients, with changes in peripheral tissues differing from those in the tumor microenvironment and taking place in phases during tumor growth. This tumor-experienced immune system mounted dampened responses to orthogonal challenges, including reduced T cell activation during viral or bacterial infection. Disruptions in T cell responses were not cell-intrinsic but rather due to reduced responses in antigen-presenting cells (APCs). Promoting APC activation was sufficient to restore T cell responses to orthogonal infection. All systemic immune changes were reversed with surgical tumor resection, revealing remarkable plasticity in the systemic immune state, which contrasts with terminal immune dysfunction in the tumor microenvironment. These results demonstrate that tumor development dynamically reshapes the composition and function of the immune macroenvironment.

## MAIN TEXT

Exploiting the mechanisms of immune activation and suppression has rapidly expanded our toolkit against cancer, leading to diverse immunotherapeutic strategies and some impressive clinical results. The efficacy of immunotherapies is currently limited, however, to select cancer types and patient subsets, begging for a more thorough understanding of the factors that govern immune responses in cancer patients. The field has garnered a robust understanding of the changes within the tumor microenvironment (TME) that subvert immune surveillance and promote tumor growth. Heterogeneous populations of immunosuppressive myeloid cells dominate many local immune landscapes, largely acting to impede cytotoxic lymphocyte activity and survival^1–4^. Intratumoral cytotoxic CD8 T cells have been the focus of the vast majority of immunomodulatory strategies in cancer therapy. However, recent studies have demonstrated that cytotoxic T cells within the TME are highly and irreversibly dysfunctional, acquiring epigenetic programs that render them incapable of normal effector functions, such as proliferation, cytokine production, and cytolysis^5^. In parallel, we and others have found that systemic immune responses are an essential component of tumor-eradicating immunity^6–10^. Consistent with these results, activated T cells in human tumors after checkpoint blockade consist of clones not observed in the tumor before the onset of therapy^11^. These findings argue that initiating a *de novo* systemic anti-tumor immune response may be essential to achieving immunotherapeutic efficacy, especially in patients who lack a strong pre-existing T cell response to their tumor.

Despite evidence that a systemic response is required for cancer rejection, our understanding of how cancer development impacts the systemic immunity remains limited. Several lines of evidence suggest systemic immune perturbations in the presence of a tumor. Peripheral granulocytic and monocytic differentiation and expansion accompanies tumor progression^12–14^, including a reduction in the number of conventional dendritic cells in bone marrow and blood^15^. Systemic effects on lymphocytes remain poorly understood. Additionally, most studies have explored anti-tumor immune responses at a single, static time point, leaving the dynamicity of the immune system during cancer development an open question. A comprehensive definition of the tumor-experienced immune macroenvironment and how it emerges over disease progression remains a crucial avenue of investigation.

A plethora of immunotherapies and vaccines seek to elicit new immune responses in cancer patients, yet no consensus has emerged for the cellular and molecular requirements to achieve this goal. In other contexts, prior immune experience has important consequences for the response to new stimuli. Chronic infection and inflammation impact immune responsiveness to novel challenges by shifting basal cytokine levels, innate immune activation states, and overall lymphocyte composition^16–18^. A detailed assessment of how tumor burden impacts responses to secondary immune challenges has yet to be performed, despite the fact that many patients likely require new immune responses to benefit from immunotherapies. It is also unclear whether there are lasting immune impacts after successful primary tumor clearance. One study suggests that the accumulation of immunosuppressive myeloid cells in the spleen rapidly alleviates with tumor resection^19^, highlighting a dynamic interaction between tumor burden and immune state. Defining the functional capacity and stability of the tumor-experienced immune macroenvironment is critical for improving immunotherapies.

The advent of high content single-cell analysis and corresponding analytical methods now allows us to tackle the challenge of characterizing systems-level immune responses in cancer. Here, we defined the systemic immune landscape in response to tumor development across eight commonly used mouse models of cancer. These data, which are now publicly available, provide a rich resource for assessing the relevance of any model to a particular question of interest or tumor type. While each tumor has unique immunological consequences, we found that three distinct models of breast cancer converged on similar changes to the systemic immune state. Tumor burden led to dynamic shifts in the organization and functional capacity of immune cells across the organism, which culminated in attenuated responses to secondary immune challenges. Tumor resection was sufficient to revert the systemic immune landscape back to a healthy baseline. These findings have implications for how and when we apply immunomodulatory agents in cancer, emphasizing the importance of strategies that are informed by alterations in the immune macroenvironment.

### Systemic immune organization is altered across multiple tumor types

We began by examining the TME across several commonly used mouse tumor models, which spanned genetically-engineered and transplantable syngeneic models across different mouse strain backgrounds. We characterized a well-established, but pre-terminal tumor stage, to reflect the patient populations most often treated with immunotherapies, but also to avoid the confounding impact of end-of-life processes. When tumors reached approximately 1 cm^3^ in volume, we harvested the tumor along with the blood, spleen, bone marrow, and tumor draining lymph node of each tumor-burdened animal and healthy control littermate. We utilized mass cytometry to quantify the abundance and activity state of immune cell subsets (Extended Data Table 1 and Extended Data Fig. 1) and performed principal component analysis (PCA) and Statistical Scaffold Mapping^6^ to visualize and assess changes in immune cell abundances.

The immune composition of the TME was distinct between tumor types, varying in the degree of both immune infiltration and diversity (Fig. 1a and Extended Data Fig. 2a). The predominant immune cell types in many tumors were tumor-associated macrophages and other CD11b^high^ myeloid subsets, particularly in the transplantable MC38 colorectal cancer and SB28 glioblastoma models. Interestingly, both transplantable LMP pancreatic cancer and genetically induced Braf^−^Pten melanoma models showed extensive eosinophil infiltration. B16-F10 syngeneic melanoma and three models of breast cancer (transplantable cell lines 4T1 and AT3, and genetically induced MMTV-PyMT) showed less relative abundance but much greater diversity in local immune cells, including B, T, and NK cell infiltration (Fig. 1a and Extended Data Fig. 2a). The unique immune profiles across tumor types are reflected by PCA (Fig. 1b).

**Fig. 1:**
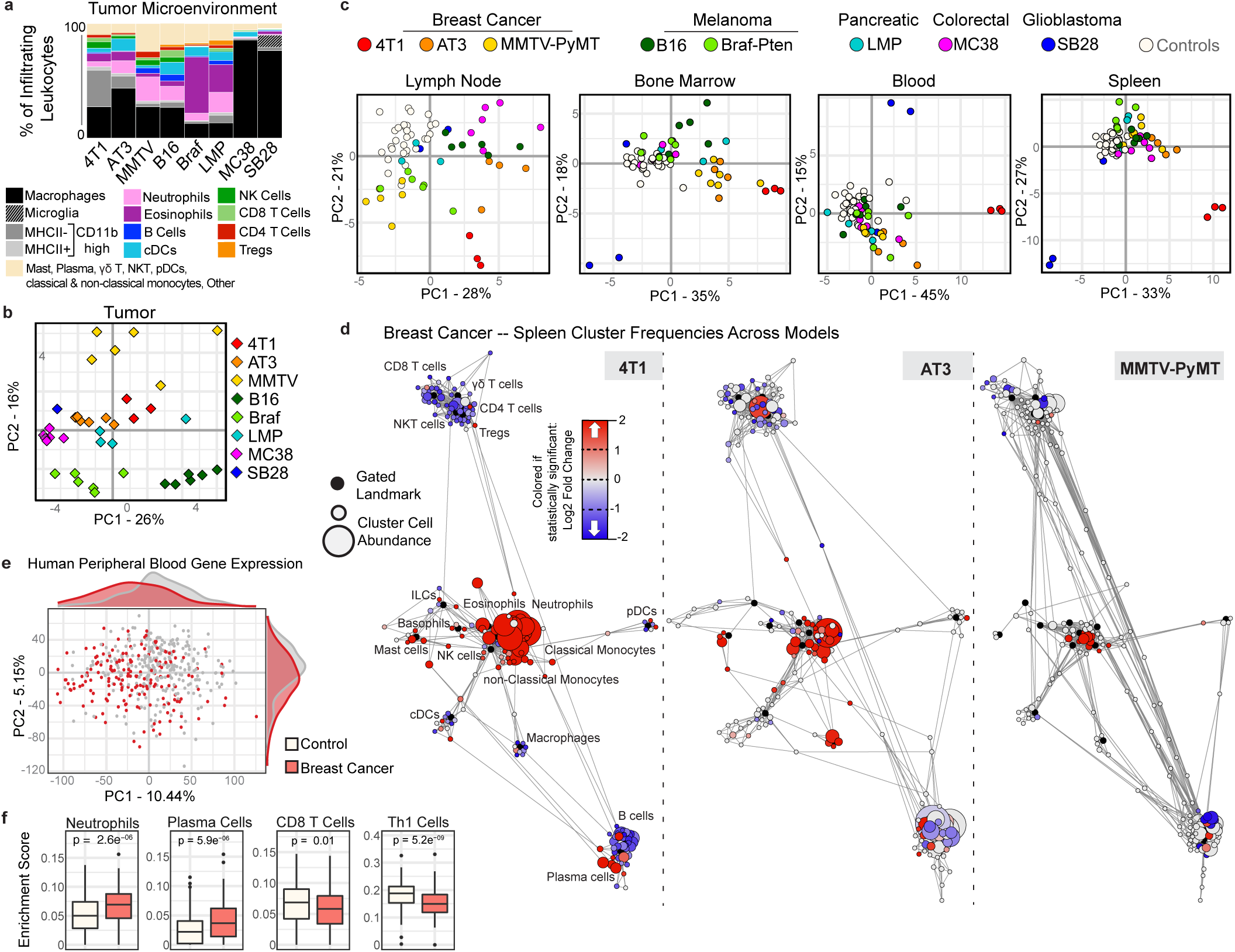
The systemic immune landscape is remodeled across tumor models. **a**, Composition of the tumor immune infiltrate across mouse tumor models at late stage tumor burden, identified manually. **b-c**, Principal component analysis (PCA) of the tumor infiltrating immune cell frequencies (**b**), and the log2 fold change of immune cell frequencies for the tumor draining lymph node, bone marrow, blood, and spleen (**c**) identified manually. **d**, Scaffold maps of spleen immune cell frequencies in breast tumor models (4T1, AT3, and MMTV-PyMT). Black nodes represent canonical cell populations identified manually. Other nodes reflect unsupervised clustering of leukocytes (see Methods). Red denotes populations significantly higher in frequency in tumor-burdened animals compared to healthy; blue denotes significantly lower frequency. For significant nodes, degree coloring reflects log2 fold change. **e-f**, PCA (**e**) and significant immune changes by cellular enrichment analysis (**f**) from human whole blood gene expression, comparing breast cancer patients (n = 173) and matched controls (n = 281).

We next asked whether different tumors also resulted in distinct systemic immune landscapes. The immune compositions of the tumor draining lymph node, bone marrow, blood, and spleen were indeed altered, albeit to varying extents, across all tumor models (Fig. 1c). While varying in magnitude, the breast cancer models consistently shifted together across principal component (PC) 2 in the lymph node and PC1 in the bone marrow, blood, and spleen. Surprisingly, SB28 glioblastoma drove extensive and distinct shifts in systemic immunity despite its localization in the CNS. Alterations in immune composition in these peripheral sites did not correspond with local immune infiltrate. Thus, tumor burden consistently drives changes in peripheral immune organization, highly dependent on the identity of the tumor and distinct from the patterns of immune infiltration in the TME.

We next performed Statistical Scaffold Analysis to interrogate the impact of tumor burden on individual immune cell types, focusing initially on the spleen as an example of a secondary lymphoid organ with immune responses initiated distal from the tumor (Fig. 1d and Extended Data Fig. 2b-f). Our approach enabled a detailed analysis of each major immune subset, building a complete picture of tumor-driven immune reorganization. All models exhibited expansions in the splenic myeloid compartment, which was dominant in some tumors, such as breast (Fig. 1d) but less dramatic in others, such as melanoma (Extended Data Fig. 2e-f). Extensive splenic remodeling in breast cancer was specifically characterized by relative increases in neutrophils, eosinophils, monocytes, and plasma cells and reductions of B and T cells (Fig. 1d). Again consistency was observed across breast cancer models, which span three mouse strain backgrounds (BALB/c for 4T1, C57BL/6 for AT3, and FVB/N for MMTV-PyMT), both orthotopic injection and spontaneous tumorigenesis, and a range of metastatic potential. Consistency despite these model differences argues strongly for a tumor and/or site-specific bias in systemic immune responses. In line with the mouse models, gene expression analysis of whole blood from untreated breast cancer patients and matched controls from the Norwegian Women and Cancer Study demonstrated a marked shift in the immune state (PC1 Wilcoxon rank sum p-value = 5.0*10^−12^, PC2 p-value = 1.6*10^−6^) (Fig. 1e). Cellular enrichment analysis demonstrated increases in neutrophils and plasma cells, as well as decreases in Th1 and CD8 T cells (Fig. 1f). Altogether, these data suggest that tumor burden broadly drives distinct immune macroenvironments, providing context to inform therapeutic manipulations designed to activate local versus systemic responses.

### Tumor growth drives non-linear changes in immune cell frequencies over time

Tumors develop gradually, yet in the clinic tumors are sampled at one point in their development to provide prognostic information related to the immune response. To understand the dynamics that result in a given local and systemic immune response, we delved further into global immune remodeling over time. Given the pronounced and consistent systemic immune changes observed in breast cancer models, we focused on these tumor settings. We began our analysis of immune cell dynamics in an orthotopic syngeneic model (4T1) due to its highly predictable kinetics before confirming results in an unrelated spontaneous model (MMTV-PyMT). We first asked whether tumor-driven immune changes developed discretely with tumor onset or progressively over tumor development. The absolute cell count of tumor-infiltrating leukocytes also positively correlated with tumor growth, supporting a progressive immune response (Extended Data Fig. 3a, r = 0.6, p = 0.0256). While absolute spleen cell counts increased along with spleen size during tumor development, cell frequencies as a percent of total leukocytes were comparable to absolute cell numbers per milligram of spleen tissue (Extended Data Fig. 3b). Thus, cell frequency was illustrated as the primary measure. Deep profiling of both the tumor and splenic immune compositions by mass cytometry revealed nonparametric correlations in individual cluster frequencies with time (Fig. 2a-b), demonstrating at the single cell level that immune changes are indeed progressive. PCA of immune cell frequencies showed progressive changes across tissues over tumor growth in both 4T1 (Fig. 2c-d) and MMTV-PyMT tumors (Extended Data Fig. 3c). Importantly, the immune profile within the TME remained distinct from those observed in peripheral sites. The draining lymph node immune composition was unique, while coordinated changes were more apparent across the spleen, blood, and bone marrow. Neutrophil expansion in the spleen and bone marrow, culminating in elevated blood circulation, but lack of accumulation within the lymph node or tumor, is one feature contributing to these unique profiles (Fig. 2d).

**Fig. 2:**
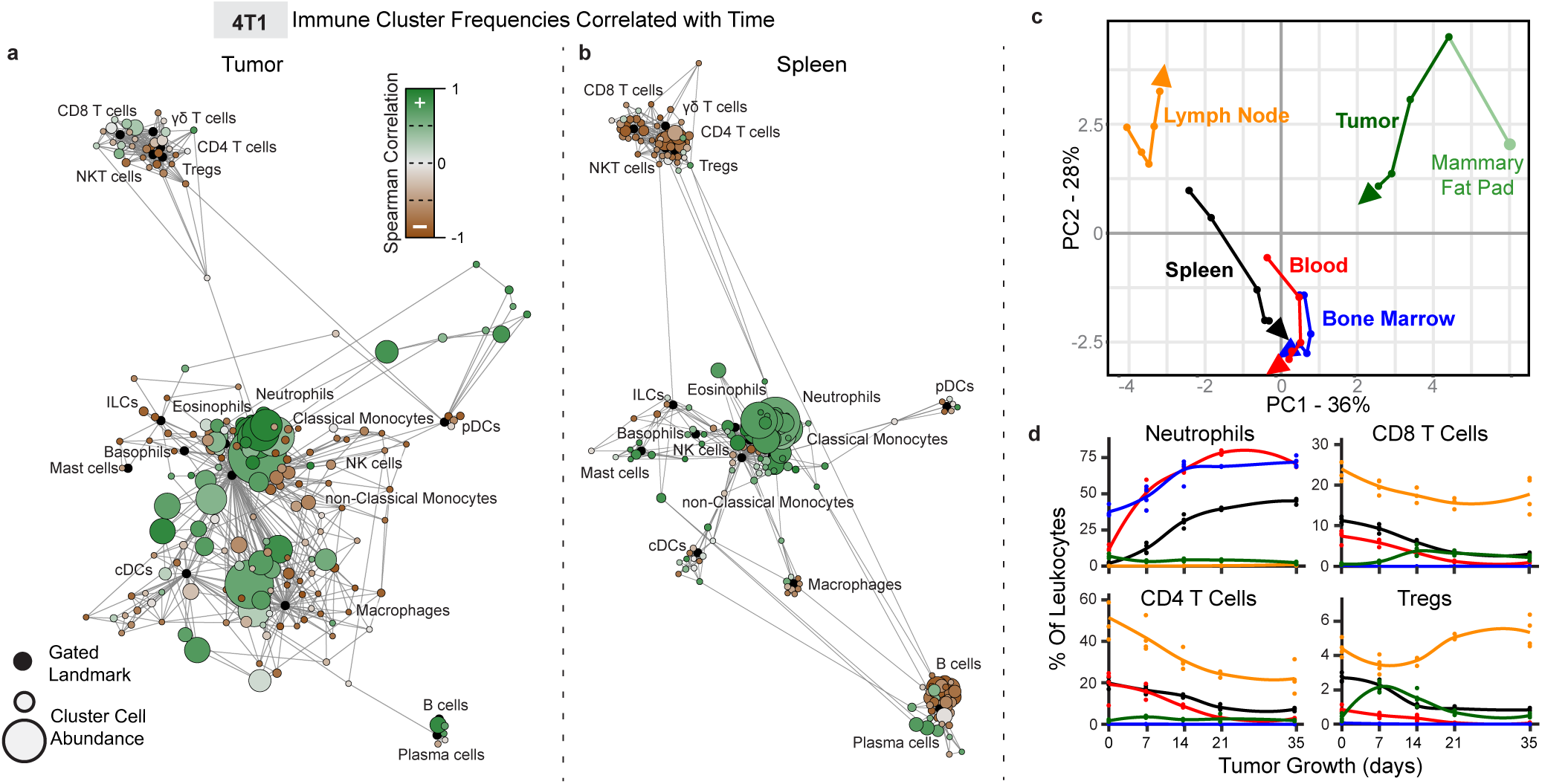
The systemic immune landscape is remodeled progressively with tumor development. **a-b**, Scaffold maps of 4T1 tumor (**a**) and spleen (**b**) cell frequencies colored by significant Spearman correlation with time (across day 0, 7, 14, 21 and 35). Green denotes positive correlation, and brown denotes negative correlation. **c**, PCA of immune cell frequencies from each immune tissue over 4T1 breast tumor growth. Vectors designate progression from control day 0 (first point) to day 7, 14, 21, and 35 (last point, arrowhead). **d**, Curves of mean cell frequencies across time from immune cell types contributing to c, colored by tissue corresponding with c.

Progressive systemic immune responses to tumor burden were not strictly linear. Rather, unique shifts in immune composition occurred at each analyzed time of tumor development. The magnitude of change was non-uniform between each time point as evident by the PCA (Fig. 2c and Extended Data Fig. 3c). While some population changes were relatively continuous, such as increasing neutrophils or decreasing CD4^+^ T cells, others were dynamic, like CD8^+^ T cells and Tregs, which reciprocally expanded and contracted at distinct times in the tumor and draining lymph node (Fig. 2d). To capture the behavior of more specific immune cell clusters over time, we constructed Statistical Scaffold maps comparing cluster abundances between each consecutive time point over 4T1 tumor growth (Extended Data Fig. 3d and Extended Data Fig. 4). In the spleen, expansion within the myeloid compartment began by day 7 and continued to day 14, preceding the relative decline in the T and B cell compartments that became evident by day 14 and continued through day (Extended Data Fig. 3d). The lymph node also showed the most dramatic immune changes by day 14 (Extended Fig. 4a), while changes in blood were more continuous (Extended Data Fig. 4b). The bone marrow and tumor contained less mature and clearly defined cell types, with many more inter-cluster connections and individualized patterns of change over tumor growth (Extended Data Fig. 4c-d). These data overwhelmingly demonstrate that the tumor immune response is a highly dynamic process.

### Immune cell states are dynamically altered across immune organs with tumor growth

We were surprised by the dramatic alterations in T cells across tissues in the periphery, as the dominant mechanisms of T cell suppression in cancer are thought to occur only in tumor antigen-specific T cells as a consequence of chronic antigen exposure^20^, or as a consequence of local immunosuppression within the TME^21^. To understand the extent of these broader systemic impacts on T cells, we leveraged unsupervised cell clustering to identify changes in T cell subsets and cell states, as well as the potential coordination of responses across organs, during tumor growth. Because immune cell frequencies are compositional, we calculated the frequencies of individual T cell clusters as a percent of total T cells in each organ to distinguish changes in T cell composition from changes in other cell types. Dramatic changes in T cell subsets were observed at specific time points, including at an intermediate stage (day 14 for 4T1, 50mm^2^ for MMTV-PyMT) and at a late stage (day 35 for 4T1, 400mm^2^ for MMTV-PyMT) (Fig. 3a, Extended Data Fig. 5a-b). Tissues contained both unique and shared T cell subsets that shifted with tumor growth (Fig. 3b-c, Extended Data Fig. 5c-e). The blood and spleen profiles were more similar and dominated by CD4^+^ T cells. In contrast, the tumor T cell pool had more shared subsets with the bone marrow, including an increasing double negative T cell population and a decreasing NKT cell population with tumor progression (Fig. 3c).

**Fig. 3:**
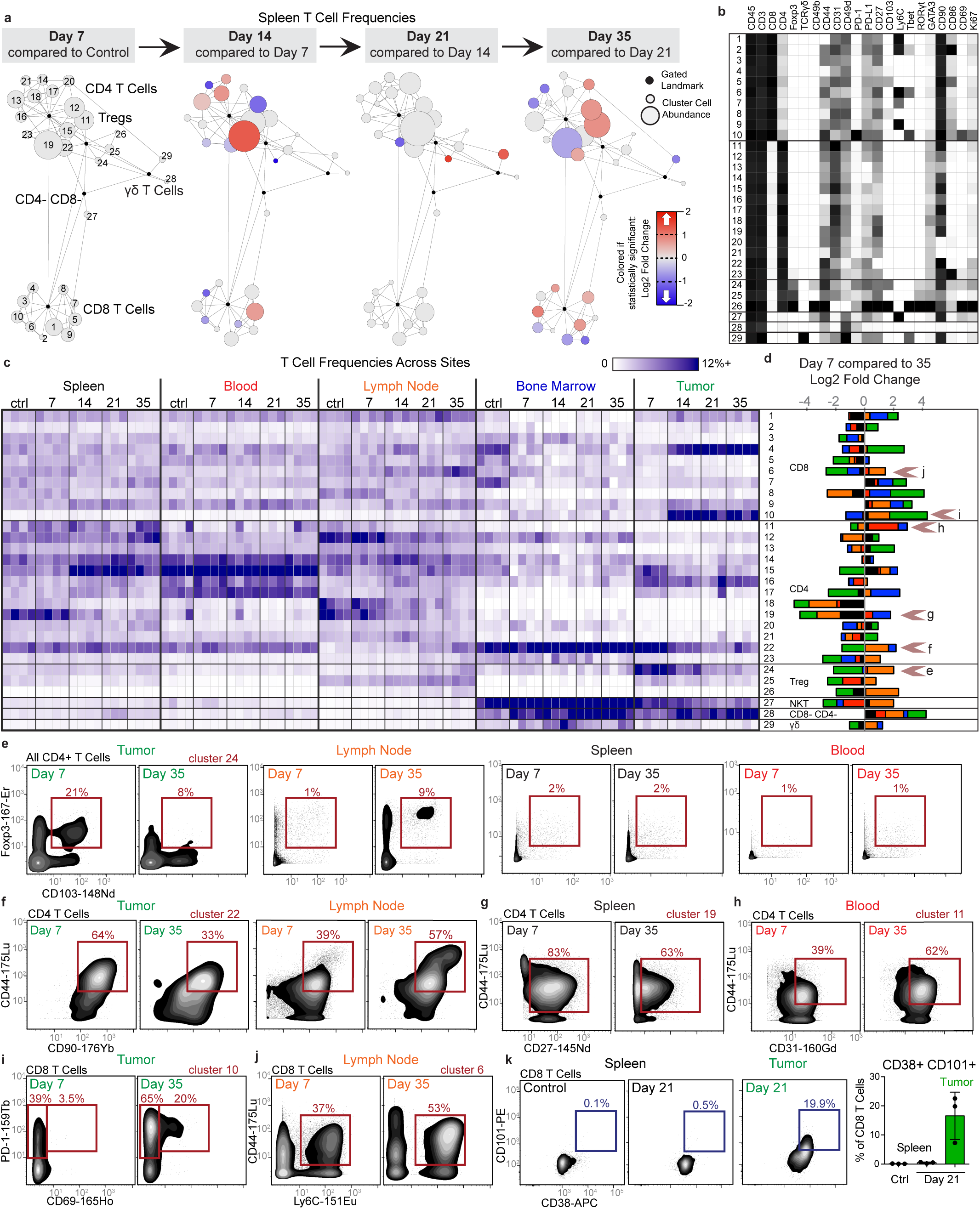
Tumor burden progressively changes the systemic T cell composition. **a-d**, CD3+ CD11b-leukocytes from all tissues clustered together from healthy and 4T1 tumor-burdened animals at progressive time points. **a**, Scaffold maps of the T cell cluster frequencies in the spleen at each disease stage, colored by fold change in frequency compared to the previous time point. **b**, Heatmap of the protein expression defining each T cell cluster, column normalized to each protein’s maximum positive expression. **c**, Heatmap of each T cell cluster frequency, by row, in each site and across the individual 3-4 animals per time point. **d**, Stacked bar plot of the log2 fold change in cluster frequency between early (day 7) and late (day 35) disease stage, colored by tissue. **e-j**, Representative scatter plots of key proteins defining T cell clusters that change in frequency in the designated tissues between early and late disease stage for Tregs (**e**), CD4 T cells (**f-h**), and CD8 T cells (**i-j**). **k**, Representative scatter plots and quantification of CD101+ CD38+ dysfunctional CD8 T cells in the spleen and tumor of health or day 21 tumor-burdened animals.

Demonstrating the breadth of immune reorganization in cancer, all T cell clusters changed in abundance across multiple tissues between early and late disease time points (Fig. 3d). Of particular interest, tumor-infiltrating CD103^+^ Tregs, described as potent suppressors of effector T cells^22^, were abundant at day 7 but decreased with tumor progression (Fig. 3e). This corresponded with CD103^+^ Treg expansion selectively in the draining lymph node, suggesting that distal suppressive mechanisms may support local changes to maintain a tumor-promoting systemic state. Anti-correlated changes extended to conventional CD4 T cells, where CD44^+^ CD90^high^ activated CD4 T cells decreased in the tumor but expanded in the lymph node (Fig. 3f). The spleen showed the greatest change in CD44^+^ CD27^+^ memory CD4^+^ T cells, which decreased with disease progression (Fig. 3g). The blood showed expansion in activated CD44^+^ CD4^+^ T cells expressing the CD31 adhesion receptor, which can promote T cell survival in settings of inflammation (Fig. 3h)^23^. CD44+ CD8+ T cells expanding in lymph node expressed Ly6C (Fig. 3j), which can support lymph node homing of central memory T cells^24^. CD8^+^ T cells generally expanded in the tumor, but the most dominant cluster expressed high levels of PD-1 and CD69 previously associated with T cell dysfunction (Fig. 3i)^25,26^. To explore the extent of dysfunction, we interrogated intratumoral and splenic T cells for their expression of CD101 and CD38, two markers recently identified as evidence of permanent T cell dysfunction^5^. Late-stage tumor burden led to accumulation of CD38^+^CD101^+^ CD8^+^ T cells in the tumor as expected; however, this phenotype did not emerge in the spleen (Fig. 3k), suggesting that CD8^+^ T cells are altered differently in the TME and in the periphery. Similar changes in T cell composition were observed in the MMTV-PyMT model (Extended Data Fig. 5c-h).

We ran a similar pan-organ clustering analysis for the mononuclear phagocyte subsets (Extended Data Fig. 6), and again found correlated and anti-correlated changes in cell states across sites with tumor progression. As expected, the tumor-infiltrating subsets were very distinct from peripheral subsets and expressed high levels of PD-L1.

We also specifically interrogated the expression dynamics of the PD-1 and PD-L1 immune checkpoint proteins, the most commonly manipulated pathway by cancer immunotherapies to facilitate T cell responses^27^. While expression of these molecules is used clinically for patient stratification, it remains unclear whether they are expressed consistently or modulated dynamically over time. We indeed found dynamic PD-1 and PD-L1 expression on infiltrating immune cells and non-immune cells of the TME (CD45^−^ CD31^−^) for both 4T1 and AT3 breast cancer models (Extended Data Fig. 7a-b). Varied expression over time held true in peripheral lymphoid organs, particularly the spleen and blood (Extended Data Fig. 7c). In fact, while the overall amount of PD-L1 expression was significantly less in the blood compared to the tumor, median leukocyte signal intensity was strongly positively correlated between these tissues (Extended Fig. 7d, r = 0.7487, p = 0.001). Both PD-1 and PD-L1 were promiscuously expressed across immune cell types, particularly within the TME (Extended Data Fig. 7e). The most prominent cells expressing PD-L1 in the periphery were non-classical monocytes^28^ and cDCs, while PD-1 was abundantly expressed on T cells, neutrophils and eosinophils. Dynamicity in PD-1 and PD-L1 expression suggests the potential for differential sensitivity to checkpoint blockade over the course of tumor development.

One potential mechanism by which immune composition could be altered is a change in cellular proliferation or death rates. By assessing Ki67 expression, we discovered that immune proliferation indeed fluctuated systemically across breast cancer models (Extended Data Fig. 8a). Changes in proliferation were highly compartmentalized such that proliferation dynamics were unique to each site but coordinated across all immune cell subsets within that site (Extended Data Fig. 8a-d). We also measured the expression of cleaved Caspase-3 to assess cell death and observed only minor changes in the spleen (26 of 200 clusters changed significantly at day 14). Changes in Ki67 and cleaved caspase-3 expression corresponded poorly with clusters that were increasing or decreasing in frequency in the spleen (Extended Data Fig. 8e). Thus, while tumor burden systemically alters proliferation and death, these processes alone likely do not account for the systemic immune alterations observed.

### *De novo* T cell responses are impaired by pre-existing malignancy

Having established that tumor development drives an altered immune macroenvironment, we determined whether immune responses to new challenges would be affected. Type 1 immune responses are associated with strong cellular immunity and are generally thought to provide optimal anti-tumor immunity. As model systems to understand how type 1 immune responses might take place in the context of cancer, we challenged healthy or AT3 tumor-burdened mice with two well-described pathogens that induce potent type 1 immunity, including CD8^+^ T cell proliferation and differentiation: lymphocytic choriomeningitis virus (LCMV) and *Listeria monocytogenes (Lm)*^29,30^. Tumor-burdened mice still cleared the pathogens from the spleen (Fig. 4a-b), consistent with the lack of complete immunosuppression in solid tumor patients. However, the cellular immune response to infection was dramatically altered. The composition of CD8 T cells was significantly altered in tumor-burdened mice after infection, with marked reductions in short-lived and memory effector CD8 T cells (Fig. 4c). CD8^+^ T cell proliferation was significantly abrogated under both infection conditions (Fig. 4d), along with impaired cytotoxic capacity indicated by a reduction in Granzyme B production (Fig. 4e). Because strong CD8^+^ T cell responses are paramount to effective anti-tumor immunity, this impairment of new cellular immunity in the context of cancer presents a fundamental and unappreciated obstacle for immunotherapy.

**Fig. 4:**
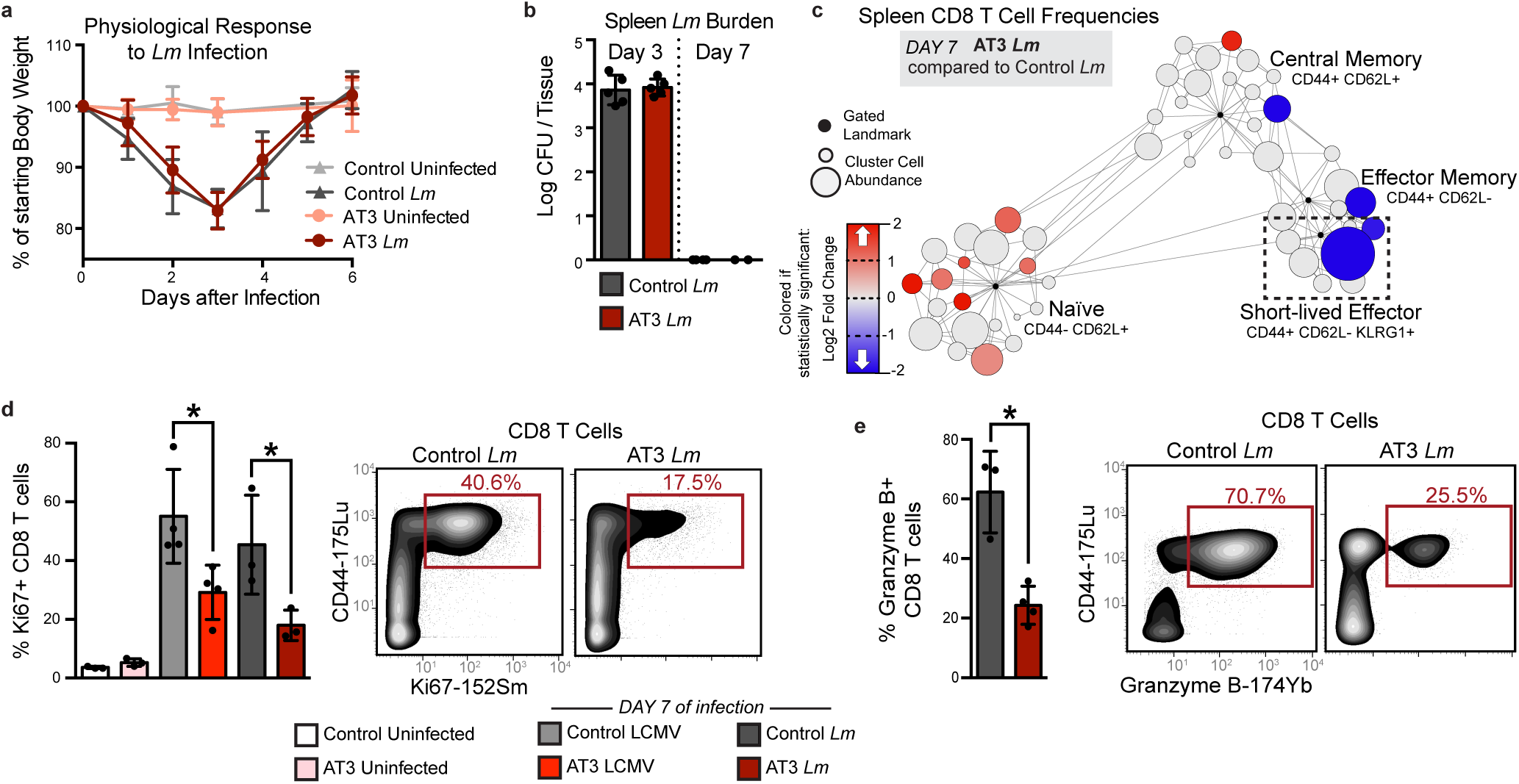
Tumor burden leads to impaired T cell responses to secondary infection. **a**-**b**, Fold change in body weight after Listeria monocytogenes (*Lm*) infection (**a**), and quantification of *Lm* bacterial burden (**b**) in control and AT3 tumor-burdened animals. **c**, Scaffold maps of CD8 T cell frequencies in the spleen in AT3 tumor-burdened mice after 7 days of *Lm* infection, colored by fold change in frequency compared to infected control mice. **d-e**, Quantification and representative scatter plots of splenic CD8+ T cell proliferation (**d**) and Granzyme B production (**e**) in response to LCMV Armstrong or *Lm* in healthy or AT3 tumor-burdened animals.

We previously found that CD8+ T cells with markers of terminal dysfunction were only observed in the TME and not the spleen (Fig. 3k). Consistent with this hypothesis, splenic CD8^+^ T cells harvested from either control or tumor-burdened animals were equally capable of producing the key effector cytokines IFNγ, TNFα, and IL-2 *in vitro* (Extended Data Fig. 9a). To test their functionality in the context of infection, CD8^+^ T cells from OT-I transgenic mice expressing a high affinity T cell receptor specific for ovalbumin (SIINFEKL) were isolated from control or tumor-burdened mice. We confirmed that AT3 tumors still drove systemic changes in TCR transgenic mice (Extended Data Fig. 9b). These cells were labeled with different fluorescent dyes to mark proliferation and were transferred together into healthy recipient mice immediately prior to infection with *Lm*-expressing ovalbumin. OT-I CD8^+^ T cells from control and tumor-burdened mice proliferated equivalently (Fig. 5a). However, when OT-I T cells were transferred into tumor-burdened recipients prior to infection, they expanded poorly, failed to induce T-bet expression associated with differentiation into effector cells, and expressed elevated levels of PD-1 (Fig. 5b). Similar results were also observed when polyclonal CD8 T cells from control or tumor-burdened mice were competitively transferred (Fig. 5c). These results demonstrate that cell extrinsic mechanisms suppress systemic T cell activation and function in the tumor context. Importantly, they also suggest that T cell behavior *in vitro* may not accurately predict their behavior once introduced into a tumor-burdened host, bearing implications for adoptive T cell therapies.

**Fig. 5:**
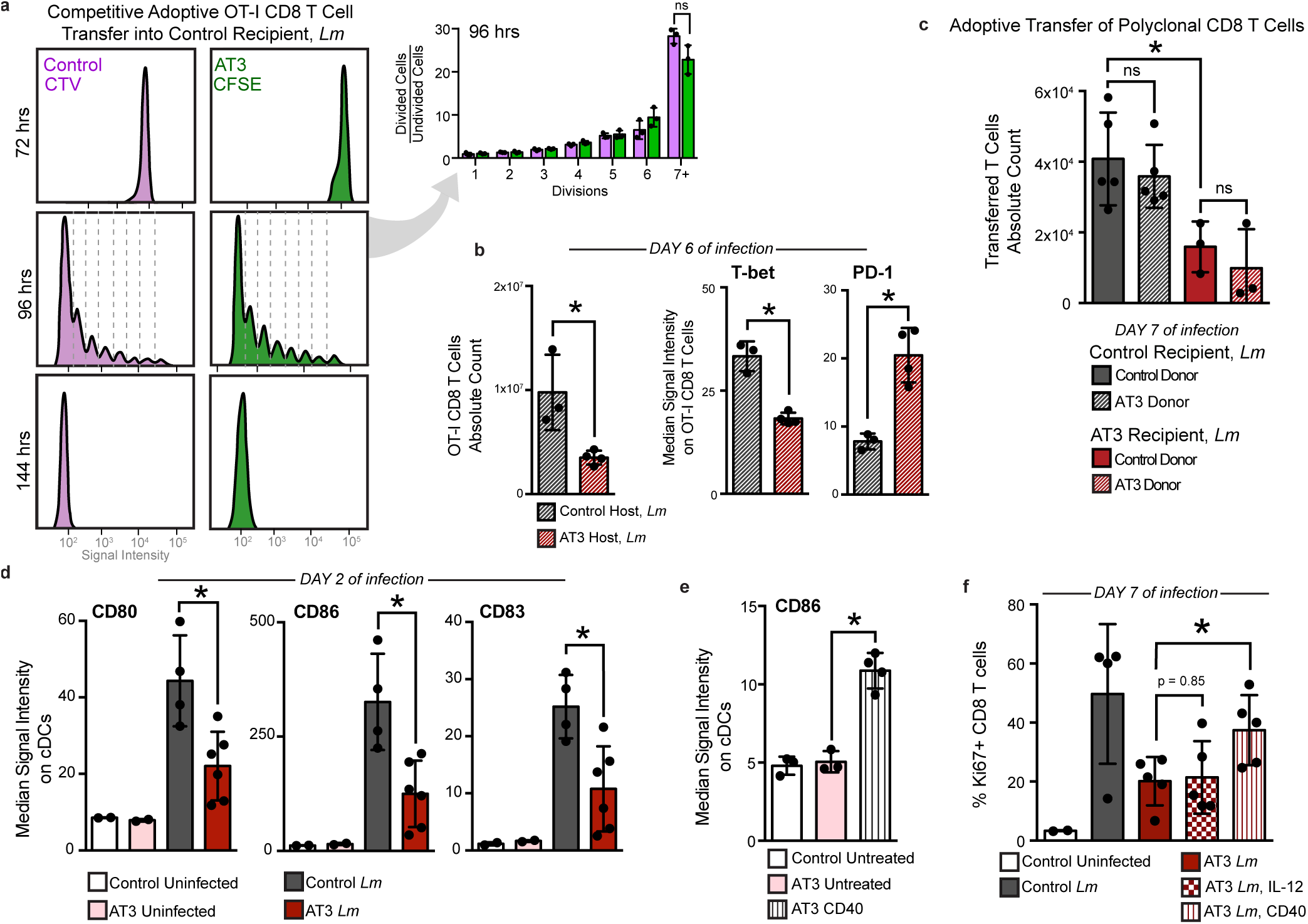
Tumor burden attenuates dendritic cell activation during secondary infection. **a**, Proliferation dyes on OT-I T cells harvested from control or tumor-burdened animals, adoptively transferred into control recipients, and analyzed at 72, 96, and 144 hours post infection with *Lm*-Ova. Quantification shown for 96 hours. Dyes diluted out by 144 hours. **b**, Absolute cell count of adoptively transferred OT-I T cells and their median signal intensity of T-bet and PD-1 at day 6 of *Lm*-OVA infection. **c**, Absolute cell count of competitively transferred polyclonal CD8 T cells from congenic (CD45.1+ AT3 tumor-burdened or CD45.1+CD45.2+ control) donors into CD45.2 control or AT3 tumor-burdened recipients, after 7 days of *Lm* infection. **d**, Median signal intensity of costimulatory proteins CD80 and CD86, and activation marker CD83 on splenic classical dendritic cells (cDCs) from healthy or AT3 tumor-burdened (day 28) mice, at day 2 of *Lm*-OVA infection. **e**, Median signal intensity of CD86 on splenic cDCs from untreated or CD40 treated AT3 tumor-burdened (day 21) mice. **f**, Quantification of splenic CD8+ T cell proliferation in response to *Lm*-OVA in healthy versus untreated, IL-12p70 treated, or CD40 treated AT3 tumor-burdened animals at day 7 of infection. p*<0.05, two-tailed t-test.

Since tumor-experienced CD8^+^ T cells in the periphery were not dysfunctional, we hypothesized that impaired APC activity earlier during the course of infection may contribute to decreased peripheral CD8^+^ T cell activation. Dendritic cells (DCs) play a key role in orchestrating CD8^+^ T cell responses to *Lm*^31^, and there is evidence to suggest that circulating DCs in breast cancer patients have reduced antigen presentation^32^. Therefore, we quantified costimulatory molecule expression on splenic DCs 2 days post infection with *Lm*. We found that DCs from AT3 tumor-burdened animals expressed lower levels of key costimulatory molecules CD80 and CD86 and the activation marker CD83 when compared to healthy controls (Fig. 5d and Extended Data Fig. 9c). At a later time point coinciding with peak T cell responses (day 7 post-infection), DCs from tumor-burdened mice continued to exhibit signs of suboptimal activation, expressing lower levels of the adhesion molecule CD54 (ICAM-1) and PD-L1 (Extended Data Fig. 9d). The latter result rules out the possibility that the PD-1/PD-L1 axis causes the impairment in T cell responses and indicates that alternative strategies are likely required to induce new systemic T cell activity. We therefore sought to pharmacologically boost APC activation as a plausible strategy for achieving this goal. Anti-CD40 treatment drives potent and systemic APC activation as shown by elevated CD86 and PD-L1 on splenic DCs (Fig. 5e and Extended Data Fig. 9e). In the context of infection, anti-CD40 treatment rescued the defect in CD8^+^ T cell proliferation in tumor-burdened animals 7 days post infection with *Lm* (Fig. 5f). At this time point, we also observed significantly higher levels of activation markers CD54 and PD-L1 on DCs after treatment (Extended Fig. 9d), consistent with enhanced APC stimulation. In stark contrast, even high doses of IL-12p70 or treatment with anti-CTLA-4 failed to rescue T cell proliferation (Fig. 5f and Extended Fig. 9f), suggesting that T cell targeted interventions alone are not sufficient. These experiments demonstrate that APCs fail to drive optimal new T cell responses in the context of tumor burden. Furthermore, these data suggest that effective immunotherapies should seek to boost APCs in combination with T cell focused treatments to fully enable *de novo* immune responses.

### Tumor resection reverses changes in systemic immune organization and responsiveness

Given that defects in T cell proliferation and differentiation were reversed when T cells were removed from a tumor-burdened context, we asked whether tumor clearance was sufficient to revert all changes in systemic immune organization and function. We performed surgical resection of tumors at a time when systemic changes were evident across sites and allowed mice to recover from surgery for an additional 14 days to mitigate immune confounders from wound healing. We carefully tracked both local recurrence and metastatic outgrowth by bioluminescent imaging. Impressively, we found that successful tumor resection reversed changes in systemic immunity in both the AT3 and 4T1 tumor models (Fig. 6a). Changes in both splenic immune cell frequencies and proliferative behavior became comparable to control animals across tissues (Fig. 6b-c, and Extended Fig. 10a-b). PCA of all major cell frequencies from both spleen and draining lymph node showed that resected animals closely resemble healthy controls along the first principal component (PC1: 43% of the variance for AT3, 57% for 4T1) (Fig. 6d). Similarly, the composition of T cell clusters in the spleen and lymph node was also largely reverted after resection (Fig. 6e). Finally, we asked whether the deficits in DC and T cell responses to infection were alleviated with tumor resection. We observed higher CD86 and PD-L1 expression on DCs at day 7 after *Lm* infection in resected mice, (Extended Fig. 10c-d) and both T cell proliferation and Granzyme B production after *Lm* infection were restored (Fig. 6f-g). Resected mice that had local or metastatic recurrence again showed deficits in DC activation and T cell responses (Extended Fig. 10c-e). Thus, changes in the systemic immune macroenvironment, unlike those of T cells in the TME, are highly dependent on the continual presence of the tumor and are dramatically reversible upon effective tumor clearance.

**Fig. 6:**
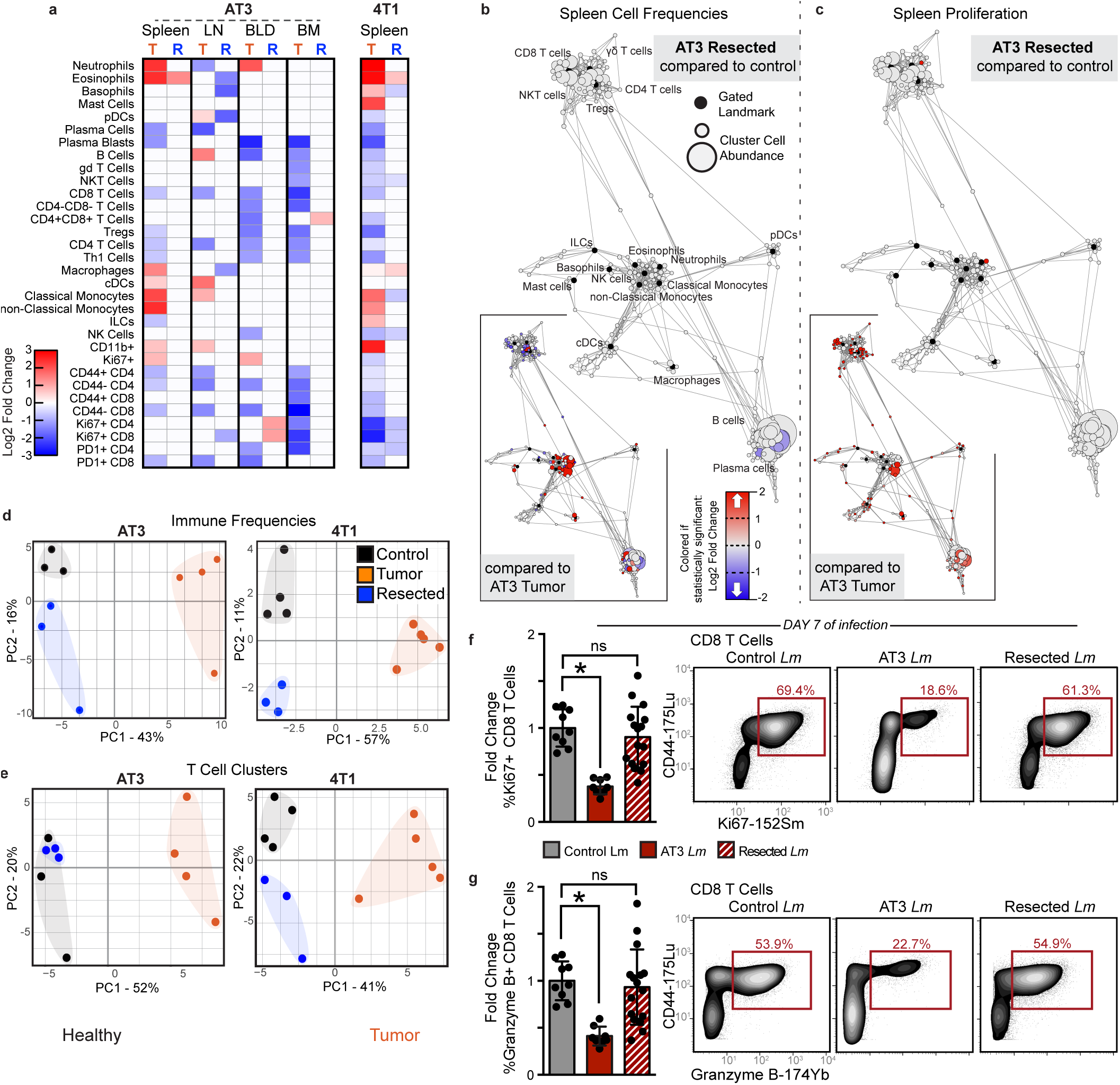
Tumor resection completely resets the systemic immune landscape. **a**, Heatmap of immune cell frequencies from tumor-burdened, T, or resected, R mice in peripheral tissues, shown as log2 fold change from control. **b-c**, Scaffold maps of spleen immune cell frequencies (**c**) and proliferation by Ki67 expression (**c**) in AT3 resected mice compared to healthy control. Insets show resected compared to tumor-burdened mice. **d-e**, PCA of all immune cell frequencies (**d**) or T cell cluster frequencies (**e**) from the spleen and tumor draining lymph node of control, tumor-burdened, or resected mice. **f-g**, Quantification and representative scatter plots of splenic CD8+ T cell proliferation (**f**) and Granzyme B production (**g**) in response to *Lm* infection in healthy, AT3 tumor-burdened, or resected mice (n = 1 to 13 per group, across 3 independent experiments). p*<0.05, two-tailed t-test.

## Discussion

This study constructs a comprehensive definition of the immune macroenvironment in cancer, capitalizing on recent technological advances to capture immune alterations across eight commonly used model systems, five sites of immune responses, and five time points in tumor progression, with 40 proteins quantified on an average of two million individual cells per animal. We greatly expanded the understanding of tumor immunity by connecting local tumor interactions with corresponding perturbations in the systemic immune state. We show that immune organization is systemically disrupted across tumor types, and that these changes are distinct from the immune effects within the local TMEs. The systemic immune impacts were unique in each tumor type and accrued nonlinearly over time, suggesting unique mechanisms of immune modulation and constant tumor-immune communication.

Immunotherapies vary in efficacy across cancer types, showing success in melanoma patients but only in a small subset of breast cancer patients^33^. Evidence of a strong pre-existing T cell response is associated with clinical benefit from currently available immunotherapies. In the remaining majority of cancer patients, it is likely that priming new immune responses will be required. Here, we show that tumor burden causes varying degrees of disruption in systemic immune state across tumor types, which is subtle in melanoma but dramatic in breast cancer. We demonstrate that severe disruptions in systemic immunity in breast cancer impair *de novo* immune responses even to highly immunogenic pathogens. Impaired new type 1 immune responses represent a fundamental, but previously unappreciated, obstacle for effective immunotherapy in patients who require priming of new T cell responses. Prior studies have connected systemic changes with relapse in breast cancer patients, showing altered immune gene signatures in uninvolved lymph nodes and blood of patients with metastatic versus non-metastatic disease^34^, and more recently that levels of circulating CD45RO- Foxp3^high^ Tregs are predictive of future relapse^35^. Vast immune disruptions argue strongly for a combinatorial immunotherapeutic approach in this context. More work needs to be done to understand the extent of systemic immune alterations across cancer patients and tumor types, and how this may inform both the likelihood of disease dissemination and the optimal therapeutic strategy.

The ability of a tumor-burdened immune system to establish *de novo* immune responses is poorly defined^36–38^, yet it is clearly essential for successful anti-tumor immunity against less immunogenic tumors. Evidence exists that human cancer patients are more susceptible to opportunistic bacterial and viral infections and also mount less effective immune responses to vaccination when compared to healthy individuals^39,40^. How much of this difference is attributable to systemic impacts of tumor burden versus the effects of common cancer therapies has remained a matter of debate. We demonstrate that immunity is indeed functionally impaired as a consequence of tumor development. The coordination of adaptive immune responses to novel challenges that did not share antigens with the tumor was significantly dampened. This striking observation challenges the idea that T cell dysfunction in cancer is limited to tumor-specific T cells and driven largely by chronic antigen presentation. Instead, our data indicate impairment in the initial coordination of a T cell response by APCs, ultimately impacting T cell proliferation and differentiation. It will be important to define the tumor-driven factors involved in failure of APCs to effectively support T cell responses across different tumor contexts.

Finally, these studies reveal remarkable plasticity in the systemic immune state. Systemic immune cells removed from the physiological context of the tumor responded normally to various challenges *in vitro* and *in vivo*. Surgical tumor resection was sufficient to revert the systemic immune landscape and function ability toward a healthy baseline. Tumor resection has previously been associated with a reduction in myeloid-derived suppressor cells^19,41^. Here, we extend these observations to characterize in depth the extent to which the systemic immune state is reversibly impacted, in both organization and in function. Influenced by the physiological immune context, immunotherapies will likely have drastically different consequences when applied pre-or post-operatively.

This study demonstrates that tumor burden drives immune programs that reach beyond local interactions. This rich data resource provides systemic immune context across all cell subsets and many tumor contexts, laying the foundation for detailed studies of specific tumor macroenvironments to match our detailed understanding of tumor microenvironments. Building a complete understanding of systems-level immunity in cancer should further our ability to drive effective and rationally designed antitumor immune responses in all cancer patients.

## METHODS

### Animals

All mice were housed in an American Association for the Accreditation of Laboratory Animal Care–accredited animal facility and maintained in specific pathogen-free conditions. Animal experiments were approved and conducted in accordance with Institutional Animal Care & Use Program protocol number AN157618. Wild-type female BALB/c, C57BL/6, and B6;129 F1 mice between 8-10 weeks old were purchased from The Jackson Laboratory and housed at our facility. 4T1 (1×10^5^ cells / 100µl) or AT3 (5×10^5^ cells / 100µl) breast cancer cells were transplanted into the fourth mammary fat pad. SB28 glioblastoma cells (1×10^5^ cells / 2µl) were transplanted into the right cerebral hemisphere by stereotactic injection. MC38 colon cancer cells (1×10^5^ cells / 100µl), B16-F10 melanoma cancer cells (1×10^5^ cells / 100µl), or LMP pancreatic cancer cells (2×10^5^ cells / 100µl) were transplanted into the subcutaneous region of the flank. Female MMTV-PyMT mice were bred at Stanford University. Tyr::CreER; Braf^V600E/+;^ Pten^lox/lox^ mice were purchased from Jackson Laboratory and housed at our facility. TCR Transgenic OT-I CD45.1 mice and heterozygous CD45.2/CD45.1 mice were bred at our facility. Animals were housed under standard SPF conditions with typical light/dark cycles and standard chow.

### Cell Lines

4T1 cells were gifted from Dr. Mary-Helen Barcellos-Hoff (UCSF). AT3 cells were gifted from Dr. Ross Levine (MSKCC). For *in vivo* experiments tracking tumor growth and recurrence after resection, we used 4T1 cells expressing mCherry-Luciferase and AT3 cells expressing GFP-Luciferase. SB28 cells, derived from a NRasV12;shp53;mPGDF transposon-induced glioma^42^, were gifted from Dr. Hideho Okada (UCSF). LMP cells, derived from the Kras^G12D/+^;LSL-Trp53^R172H/+^;Pdx-1-Cre model of pancreatic cancer^43^, were gifted from Dr. Edgar Engleman (Stanford University). MC38 cells and B16-F10 cells gifted from Dr. Jeffrey Bluestone (UCSF). 4T1, MC38, B16 and SB28 cells were cultured in RPMI-1640, and AT3 and LMP cells were cultured in DMEM, all supplemented with 10% FCS, 2 mM L-glutamine,100 U/mL penicillin and 100 mg/mL penicillin/streptomycin.

### Infectious Agents

*Listeria monocytogenes* strain 10403s expressing OVA (*Lm*-OVA) was originally from Hao Shen ^44^ and kindly provided by Shomyseh Sanjabi (UCSF). Lm-OVA stocks frozen at −80 C were grown overnight at 37 C in BHI broth supplemented with 5 ug/ml Erythromycin. Then, overnight cultures were sub-cultured by diluting into fresh BHI broth supplemented with 5 ug/ml Erythromycin and grown for 4 hours. Bacteria CFU was then quantified by measuring optical density at 600 nm. Bacteria were then diluted to 5×10^4^ CFU / 100µl in sterile PBS and 100 µl was injected per mouse i.v. via the retro-orbital vein.

Lymphocytic choriomeningitis virus (LCMV) was kindly provided by Dr. Jason Cyster (UCSF) and mice were infected with pre-titered and aliquoted stocks stored in PBS at −80C and diluted with sterile PBS. Mice were infected with 2×10^5^ PFU by intraperitoneal injection.

### Mass Cytometry Antibodies

All mass cytometry antibodies and concentrations used for analysis can be found in Table S1. Primary conjugates of mass cytometry antibodies were prepared using the MaxPAR antibody conjugation kit (Fluidigm) according to the manufacturer’s recommended protocol. Following labeling, antibodies were diluted in Candor PBS Antibody Stabilization solution (Candor Bioscience GmbH, Wangen, Germany) supplemented with 0.02% NaN3 to between 0.1 and 0.3 mg/mL and stored long-term at 4°C. Each antibody clone and lot was titrated to optimal staining concentrations using primary murine samples.

### Cell Preparation

All tissue preparations were performed simultaneously from each individual mouse, as previously reported^6^. After euthanasia by C02 inhalation, peripheral blood was collected via the posterior vena cava prior to perfusion of the animal and transferred into sodium heparin-coated vacuum tubes prior to dilution in PBS with 5mM EDTA and 0.5% BSA (PBS/EDTA/BSA). Spleens and lymph nodes were homogenized in PBS/EDTA at 4°C. Bone marrow was flushed from femur and re-suspended in PBS/EDTA at 4°C. Tumors were finely minced and digested in RPMI-1640 with 4 mg/ml collagenase IV, and 0.1 mg/ml DNase I. After digestion, re-suspended cells were quenched with PBS/EDTA at 4°C. All tissues were washed with PBS/EDTA and re-suspended 1:1 with PBS/EDTA and 100mM Cisplatin (Enzo Life Sciences, Farmingdale, NY) for 60 s before quenching 1:1 with PBS/EDTA/BSA to determine viability as previously described^45^. Cells were centrifuged at 500 g for 5 min at 4°C and re-suspended in PBS/EDTA/BSA at a density between 1-10*106 cells/ml. Suspensions were fixed for 10 min at RT using 1.6% PFA and frozen at −80°C.

### Mass-Tag Cellular Barcoding

Mass-tag cellular barcoding was performed as previously described^46^. Briefly, 1*10^6^ cells from each animal were barcoded with distinct combinations of stable Pd isotopes in 0.02% saponin in PBS. Samples from any given tissue from each mouse per experiment group were barcoded together. Cells were washed once with cell staining media (PBS with 0.5% BSA and 0.02% NaN3), and once with 1X PBS, and pooled into a single FACS tube (BD Biosciences). After data collection, each condition was deconvoluted using a single-cell debarcoding algorithm^46^.

### Mass Cytometry Staining and Measurement

Cells were resuspended in cell staining media (PBS with 0.5% BSA and 0.02% NaN3) and metal-labeled antibodies against CD16/32 were added at 20 mg/ml for 5 min at RT on a shaker to block Fc receptors. Surface marker antibodies were then added, yielding 500 uL final reaction volumes and stained for 30 min at RT on a shaker. Following staining, cells were washed 2 times with cell staining media, then permeabilized with methanol for at 10 min at 4°C. Cells were then washed twice in cell staining media to remove remaining methanol, and stained with intracellular antibodies in 500 mL for 30 min at RT on a shaker. Cells were washed twice in cell staining media and then stained with 1mL of 1:4000 191/193Ir DNA intercalator (Fluidigm) diluted in PBS with 1.6% PFA overnight. Cells were then washed once with cell staining media and then two times with double-deionized (dd)H20. Care was taken to assure buffers preceding analysis were not contaminated with metals in the mass range above 100 Da. Mass cytometry samples were diluted in ddH2O containing bead standards (see below) to approximately 10^6^ cells per mL and then analyzed on a CyTOF 2 mass cytometer (Fluidigm) equilibrated with ddH2O. We analyzed 1-5*10^5^ cells per animal, per tissue, per time point, consistent with generally accepted practices in the field.

### Mass Cytometry Bead Standard Data Normalization

Data normalization was performed as previously described^6^. Briefly, just before analysis, the stained and intercalated cell pellet was resuspended in freshly prepared ddH2O containing the bead standard at a concentration ranging between 1 and 2*10^4^ beads/ml. The mixture of beads and cells were filtered through a filter cap FACS tubes (BD Biosciences) before analysis. All mass cytometry files were normalized together using the mass cytometry data normalization algorithm^47^, which uses the intensity values of a sliding window of these bead standards to correct for instrument fluctuations over time and between samples.

### Mass Cytometry Gating Strategy

After normalization and debarcoding of files, singlets were gated by Event Length and DNA. Live cells were identified by Cisplatin negative cells. All positive and negative populations and antibody staining concentrations were determined by titration on positive and negative control cell populations.

### Scaffold Map Generation

Statistical scaffold maps were generated using the open source Statistical Scaffold R package available at github.com/SpitzerLab/statisticalScaffold with modifications detailed below.

As previously described^6^, cells from each tissue for all animals were clustered together and then deconvolved into their respective samples. Cluster frequencies or the Boolean expression of specific proteins for each cluster were passed into the Significance Across Microarrays algorithm^48,49^, and the fold change results were reported (rather than the binary significance cutoff as originally implemented in Spitzer et al., 2017). Cluster frequencies were also correlated with the time from tumor inoculation using Spearman’s rank-ordered correlation. All results were tabulated into the Scaffold map files for visualization through the graphical user interface, with coloring modifications to graph the spectrum of fold change or correlation strength. The fold change was log2 normalized and graphed with an upper and lower limit of a four-fold difference, unless otherwise indicated. Cluster frequencies were calculated as a percent of total live nucleated cells (excluding erythrocytes). The spleen data from the 4T1 model was used to spatialize the initial Scaffold map because all major, mature immune cell populations are present in that tissue.

### Cell Frequency Heat Map Generation

Specified subsets, i.e. T cells and mononuclear phagocytes, were manually gated from each tissue for all animals and clustered together. Cluster frequencies were calculated as a percent of total live nucleated cells within that subset (excluding erythrocytes). T cells were identified as CD3^+^, CD11b^−^. Mononuclear phagocytes were defined as CD11b^+^, CD19^−^, CD3^−^, Ly6G^−^. Heatmaps of the resulting cluster frequencies were generated in R.

### Human Gene Expression Analysis

Whole blood microarray data was generated by The Norwegian Women and Cancer (NOWAC) study and is deposited in the European Genome-Phenome Archive under accession number EGAS00001001804 as previously reported ^50^. Principal component analysis of centered and scaled data was performed in R using the prcomp function. xCell cell type enrichment analysis was performed in R using the xCell package (https://github.com/dviraran/xCell) using a customized list of cell populations known to exist in peripheral whole blood (B-cells, Basophils, CD4^+^ memory T-cells, CD4^+^ naive T-cells, CD4^+^ T-cells, CD4^+^ Tcm, CD4^+^ Tem, CD8^+^ naive T-cells, CD8^+^ T-cells, CD8^+^ Tcm, CD8^+^ Tem, cDC, Class-switched memory B-cells, Eosinophils, Erythrocytes, Megakaryocytes, Memory B-cells, Monocytes, naive B-cells, Neutrophils, NK cells, NKT, pDC, Plasma cells, Platelets, Tgd cells, Th1 cells, Th2 cells, Tregs).

### *In vitro* CD8 T cell Differentiation and cytokine production

Mice bearing 21-day AT3 tumors were euthanized and their spleens harvested and dissociated. CD8 T cells were enriched using the EasySep Streptavidin Negative Selection Kit with the following biotinylated markers: CD11b, MHCII, CD11c, Gr1, B220, CD4, CD44, and Ter119. Isolated CD8 T cells were then stimulated with plate-bound CD3 and suspended in CD28 containing T cell media for 3 days. The cells were then removed from CD3/CD28 stimulation and rested for 1 day. Cells were then restimulated with PMA & Ionomycin or left unstimulated for 4 hours with Brefeldin A and analyzed by flow cytometry.

### Adoptive T Cell Transfer

For OT1 and polyclonal adoptive transfers, CD8 T cells were isolated from spleens of CD45.1 OT1 TCR transgenic or CD45.1/CD45.2 heterozygote wildtype or CD45.1 BoyJ mice by enrichment with EasySep Streptavidin Negative Selection Kit with the following biotinylated markers: CD11b, MHCII, CD11c, Gr1, B220, CD4, and Ter119. Cells were stained with CFSE or Cell Trace Violet and 1×10^5^ cells were then adoptively transferred into each recipient mouse via the retroorbital vein.

### Quantifying Bacterial Burden

To quantify bacterial burden, spleens were harvested and dissociated. Cells from each mouse were lysed in 0.5% TritonX 100 in PBS and cells were serially diluted in duplicate and aliquots were then added to BHI agar and incubated overnight at 37C. Colonies grown were then counted to quantify bacterial CFU present.

### Treatments

All *in vivo* antibody treatments were given i.p. starting on day 0 of *Lm*-Ova infection: 200 μg of agonistic CD40 (FGK4.5, BioXCell) on day 0, 225 μg of recombinant IL-12p70 (BioLegend) daily, and 200 μg of antagonistic CTLA-4 (9H10, BioXCell) on day 0 and day 3.

### Tumor Resection

Mice bearing 14-day 4T1 tumors or 16 to 21-day AT3 tumors (between 350-550mm^3^) were anesthetized by intraperitoneal (i.p) injection with a mixture of ketamine and xylazine, and titrated to effect with isoflurane from a precision vaporizer. The surgical site was shaved and sterilized with 70% ethanol and 10% povidone iodine. An incision was made subcutaneously at the anterior midline and along the flank of the side with the tumor, using surgical scissors, to reveal the inguinal mammary tumor. The tumor was teased away using forceps and the surgical wound closed with wound clips. Wound clips were removed after 7 days. 10-20% of resected mice had tumor recurrence due to incomplete removal of primary tumors or outgrowth of micro-metastases. These mice were excluded from the experiments to which they were initially assigned.

### Flow Cytometry

Cells were stained for viability with Zombie-NIR stain. Cell surface staining was performed in cell staining media (PBS with 0.5% BSA and 0.02% NaN3) for 15 minutes at room temperature. Intracellular staining was performed after fixing cells with BioLegend FluoroFix Buffer and permeabilizing cells with Biolegend’s Intracellular Staining Perm Wash Buffer. The following anti-mouse antibodies were used: (PE-Dazzle594) – CD3 (clone 17A2), (PacificBlue) – CD4 (clone RM4-5), (BV786) – CD8 (clone 53-6.7), (APC-Cy7) – CD45 (clone 30-F11), (APC) – CD38 (clone 90), (PE) – CD101 (clone Moushi101), (PD1) – PE-Cy7 (clone 29F.1A12), (BV421) – TCRb (clone H57-597), (PE) – IFNg (clone XMG1.2), (BV711) – IL2 (clone JES6-5H4), (FITC) – TNFalpha (clone MP6-XT22), (BV650) – CD8 (clone 53-6.7), (PE) – CD45.1 (clone A20). All antibodies were purchased from Biolegend, Inc., BD Biosciences, or Thermo Fisher Scientific. Stained cells were analyzed with a CytoFLEX flow cytometer (Beckman Coulter) or an LSR II flow cytometer (BD Biosciences).

Singlets were gated by FSC-A and FSC-W, as well as by SSC-A and SSC-W. All positive and negative populations were determined by staining on positive and negative control populations.

### QUANTIFICATION AND STATISTICAL ANALYSIS

Comparison of cell frequencies and protein expression in Statistical Scaffold was performed using Significance Analysis of Micro-arrays as described above and in Bair and Tibshirani, 2004 and Bruggner et al., 2014. Analysis of principle components for human gene expression was performed using two-tailed Wilcoxon rank-Sum test in R. All comparisons over 4T1 tumor growth were performed by One-way ANOVA with Tukey correction in Prism. All other comparisons after infection, treatment, or resection were made using two-tailed t tests in Prism. All tests with q < 0.05 were considered statistically significant. Unless otherwise stated, n = 3 to 6 independent mice for each experimental condition.

### DATA AND SOFTWARE AVAILABILITY

The updated Statistical Scaffold package and all mass cytometry data will be made publicly available concurrent with publication of the manuscript.

## Supporting information

Supplementary Figures for Allen and Hiam et al.

## ACKNOWLEDGMENTS

We thank the UCSF Flow Cytometry Core and Stanley Tamaki for CyTOF maintenance, Drs. Mary-Helen Barcellos-Hoff, Ross Levine, Hideho Okada, Edgar Engleman and Jeffrey Bluestone for cell lines, transgenic mice and reagents, and Iliana Tenvooren and Diana Marquez for assistance in animal work. This work was supported by NIH grants DP5OD023056 and P50CA097257 (UCSF Brain Tumor SPORE Developmental Research Program) and investigator funding from the Parker Institute for Cancer Immunotherapy to M.H.S., and by NIH grant S10OD018040, which enabled procurement of the mass cytometer used in this study.

M.H.S. receives research funding from Roche/Genentech and Valitor Inc. and has been a paid consultant for Five Prime Therapeutics and Ono Pharmaceutical.

## AUTHOR CONTRIBUTIONS

Conceptualization, B.M.A, K.J.H., Y.C., and M.H.S.; Experimental Methodology, B.M.A, K.J.H., C.E.B., A.V., R.B., Y.C., and M.H.S.; Computational Methodology, B.M.A, and M.H.S.; Investigation, all authors; Writing – Original Draft, B.M.A.; Writing – Review & Editing, all authors; Funding Acquisition, M.H.S.; Supervision, M.H.S.

## EXTENDED DATA

*Extended Data Fig. 1: Main Mass Cytometry Gating Scheme*.

**a**, Main gating strategy for identifying major immune cell populations from mass cytometry datasets.

***Extended Data Fig. 2: Systemic immunity is distinctly remodeled across tumor models***.

**a**, Relative abundance of total leukocytes infiltrating the TME across eight tumor models. **b-f**, Scaffold maps of spleen cell frequencies across five distinct tumor models, SB28 glioblastoma (**b**), MC38 colorectal (**c**), LMP pancreatic (**d**), B16 melanoma (**e**), and Braf-PTEN melanoma (**f**), comparing late stage tumor burden to their respective health littermate controls.

***Extended Data Fig. 3: Systemic immunity is distinctly remodeled over tumor development***.

**a**, Pearson correlation between tumor mass and absolute number of infiltrating leukocytes in 4T1 breast tumors. **b**, Spleen immune absolute cell counts, adjusted absolute cell counts per mg of tissue, and unadjusted immune frequencies at each time point for neutrophils, B cells and T cells of the 4T1 breast tumor model. **c**, PCA of relative immune cell frequencies from each major immune tissue over time in the MMTV-PyMT breast tumor model. Vectors designate progression from control (first point) to 25 mm^2^, 50mm^2^, 125mm^2^, and 400mm^2^ (last point, arrowhead). **d**, Scaffold maps of immune cell frequencies in the spleen at each time point of 4T1 tumor burden, colored by log2 fold change in frequency compared to the previous time point.

***Extended Data Fig. 4: Immunity is distinctly remodeled by compartment over tumor development***.

**a-d**, Scaffold maps of immune cell frequencies over 4T1 tumor progression in the tumor draining lymph node (**a**) blood (**b**), bone marrow (**c**), and tumor (**d**), colored by fold change relative to the previous time point.

***Extended Data Fig. 5: Tumor growth shifts the systemic T cell composition across models***.

**a-b**, PCA of T cell cluster frequencies across lymphoid tissues over tumor development for the 4T1 (**a**) and MMTV-PyMT (**b**) breast tumor models. Vectors designate directional progression from control (first point) to late stage disease (last point, arrowhead). In **a**, tumor time points include day 7, 14, 21, and 35 after 4T1 cancer cell transplant. In **b**, tumor time points include tumor sizes of 25 mm^2^, 50 mm^2^, 125 mm^2^, and 400 mm^2^. **c-e**, CD3+ CD11b-leukocytes from all tissues clustered together from healthy and MMTV-PyMT tumor-burdened animals at progressive tumor sizes. **c**, Heatmap of each T cell cluster frequency, by row, in each site and across the individual 2-3 animals per time point. **d**, Stacked bar plot of the log2 fold change in cluster frequency between early (25 mm^2^) and late (400 mm^2^) disease time points, colored by tissue. **e**, Heatmap of the protein expression defining each T cell cluster, column normalized to each protein’s maximum positive expression. **f-h**, Representative scatter plots of key proteins that define T cell clusters changing in frequency in the designated site between early and late disease stage for CD8 T cells (**f**), Tregs (**g**), and CD4 T Cells (**h**).

***Extended Data Fig. 6: Tumor growth shifts the systemic mononuclear phagocyte composition***.

**a**, CD3-CD19-leukocytes from all tissues clustered together from healthy and 4T1 tumor-burdened animals at progressive time points. *Left*, stacked bar plot of the log2 fold change in cluster frequency between early (day 7) and late (day 35) times points, colored by tissue. *Right*, heatmap of the protein expression defining each cluster, column normalized to each protein’s maximum positive expression. **b**, Curves of the mean cell frequencies over time in the 4T1 breast tumor model from designated mononuclear phagocyte cell types, colored by tissue. **c**, PCA of the mononuclear phagocyte cell frequencies from each tissue over time in the 4T1 breast tumor model. Vectors designate progression from control (first point) to day 7, 14, 21, and 35 (last point, arrowhead). Coloring of tissues for a-c corresponds to labels in c.

***Extended Data Fig. 7: PD-1 and PD-L1 expression is dynamic over tumor growth***.

**a**, Distribution of PD-1 and PD-L1 signal intensities on tumor infiltrating leukocytes over time in the 4T1 or AT3 breast tumor models. Coloring of time points for a-d corresponds to legend in a. **b**, Percent of total infiltrating leukocytes (*left of dashed line*) or CD45-, non-endothelial cells (*right of dashed line*) with high PD-1 or PD-L1 expression in the 4T1 or AT3 tumor models. **c**, Percent of leukocytes with high PD-1 or PD-L1 expression over time and across tissues, 4T1 model. **d**, Pearson correlation between median PD-L1 signal intensity on blood versus tumor infiltrating leukocytes, 4T1 model. **e**, Percent of each major immune cell subset expressing high PD-1 or PD-L1 in the tumor, blood, and spleen, identified manually. Cell subsets below 0.2% of total leukocytes were not included, X. Bars ordered by time point, beginning at healthy control. Double positive PD-1/PD-L1 expression was rare and not illustrated. p*< 0.05, One-Way ANOVA, with Tukey correction versus control tissue or healthy mammary fat pad (blue in b-c, fill corresponding to bar color in e), or versus day 7 (green in b-c).

***Extended Data Fig. 8: Tumor burden induces tissue-specific changes in immune cell cycling***.

**a-b**, Log2 fold change in bulk Ki67 expressing leukocytes in each tissue tissues for 4T1, AT3 and MMTV breast tumors (**a**), and over 4T1 tumor progression (**b**). p*< 0.05, One-Way ANOVA, with Tukey correction versus control. **c-d**, Statistical Scaffold maps of Ki67 expression in immune cells of the tumor draining lymph node comparing control to day 21 (**c**) and the Spleen over time (**d**) in 4T1 tumor burdened animals. **e**, Percent of increasing clusters (red, total of 56) or decreasing clusters (blue, total of 90) that have corresponding changes in cell cycle markers Ki67 and cleaved Caspase-3.

***Extended Data Fig. 9: Tumor driven deficits in T cell responses are cell-extrinsic***.

**a**, Expression of inflammatory cytokines, INFy, IL-2, and TNFa in splenic CD8 T Cells isolated from control or AT3 tumor-burdened mice after *in vitro* differentiation with CD3, CD28 and IL-2, and re-stimulation with BrefeldinA and PMA Ionomycin. **b**, Scatter plots of CD11b and Ly6G showing expected neutrophilia in OT-I TCR transgenic mice with AT3 tumor burden. **c**, Histograms of CD80, CD86, and CD83 signal intensity on cDCs from healthy or AT3 tumor-burdened mice at day 2 of *Lm*-OVA infection. **d**, Median signal intensity of PD-L1 and CD54 activation markers on splenic cDCs from healthy or AT3 tumor-burdened mice compared to IL-12p70 or CD40 treatment at day 7 of *Lm*-OVA infection. **e**, Median signal intensity of PD-L1 on splenic cDCs from untreated or CD40 treated AT3 tumor-burdened (day 21) mice. **f**, Quantification of splenic CD8+ T cell proliferation in healthy, untreated or CTLA-4 treated AT3 tumor-burdened animals in response to 7 days of *Lm*-OVA infection. p*<0.05, two-tailed t-test.

***Extended Data Fig. 10: Tumor resection resets systemic immune organization and function***.

**a-b**, Statistical scaffold maps of spleen immune cell frequencies (**a**) and proliferation by Ki67 expression (**b**) in 4T1 resected mice compared to health control. Insets show resected mice compared to tumor-burdened mice. **c-d**, Median signal intensity of CD86 (**c**) and PD-L1 (**d**) on splenic cDCs from healthy, AT3 tumor-burdened, resected, or resected mice with recurrence at day 7 of *Lm*-OVA infection. p*<0.05, two-tailed t-test. **e**, Quantification of splenic CD8+ T cell proliferation and Granzyme B production in response to *Lm*-OVA in healthy versus resected mice with local or metastatic recurrence.

## REFERENCES

1. Azizi, E. et al. Single-Cell Map of Diverse Immune Phenotypes in the Breast Tumor Microenvironment. Cell 174, 1293–1308 (2018).

2. Joyce, J. A. & Fearon, D. T. T cell exclusion, immune privilege, and the tumor microenvironment. Science. 348, 74–80 (2015).

3. Wagner, J. et al. A Single-Cell Atlas of the Tumor and Immune Ecosystem of Human Breast Cancer. Cell 177, 1–16 (2019).

4. Tsujikawa, T. et al. Quantitative Multiplex Immunohistochemistry Reveals Myeloid-Inflamed Tumor-Immune Complexity Associated with Poor Prognosis. Cell Rep. 19, 203–217 (2017).

5. Philip, M. et al. Chromatin states define tumour-specific T cell dysfunction and reprogramming. Nature 545, 452–456 (2017).

6. Spitzer, M. H. et al. Systemic Immunity Is Required for Effective Cancer Immunotherapy. Cell 168, 487–502 (2017).

7. Fransen, M. F. & Van Hall, T. Tumor-draining lymph nodes are pivotal in PD-1/PD-L1 checkpoint therapy. JCI Insight 3, e124507 (2018).

8. Tang, H. et al. PD-L1 on host cells is essential for PD-L1 blockade-mediated tumor regression. J Clin Invest 128, 580–588 (2018).

9. Chamoto, K. et al. Mitochondrial activation chemicals synergize with surface receptor PD-1 blockade for T cell-dependent antitumor activity. PNAS 114, E761–E770 (2017).

10. Mathios, D. et al. Anti – PD-1 antitumor immunity is enhanced by local and abrogated by systemic chemotherapy in GBM. Sci. Transl. Med. 8, 1–12 (2016).

11. Yost, K. E. et al. Clonal replacement of tumor-specific T cells following PD-1 blockade. Nat. Med. 25, 1251–1259 (2019).

12. McAllister, S. S. & Weinberg, R. A. The tumour-induced systemic environment as a critical regulator of cancer progression and metastasis. Nat. Cell Biol. 16, 717–727 (2014).

13. Zhang, S. et al. The Role of Myeloid-Derived Suppressor Cells in Patients with Solid Tumors: A Meta-Analysis. PLoS One 11, e0164514 (2016).

14. Casbon, A.-J. et al. Invasive breast cancer reprograms early myeloid differentiation in the bone marrow to generate immunosuppressive neutrophils. Proc. Natl. Acad. Sci. U. S. A. 112, E566–75 (2015).

15. Meyer, M. A. et al. Breast and pancreatic cancer interrupt IRF8-dependent dendritic cell development to overcome immune surveillance. Nat. Commun. 9, 1–19 (2018).

16. Barnstorf, I. et al. Chronic virus infection compromises memory bystander T cell function in an IL-6/ STAT1-dependent manner. J. Exp. Med 216, 571–586 (2019).

17. Snell, L. M. et al. CD8 + T Cell Priming in Established Chronic Viral Infection Preferentially Directs Differentiation of Memory-like Cells for Sustained Immunity. Immunity 49, (2018).

18. Osborne, L. C. et al. Virus-helminth coinfection reveals a microbiota-independent mechanism of immunomodulation. Science. 345, 578–582 (2014).

19. Ghochikyan, A. et al. Primary 4T1 tumor resection provides critical “window of opportunity” for immunotherapy. Clin Exp Metastasis 31, 185–198 (2014).

20. Schietinger, A. et al. Tumor-Specific T Cell Dysfunction Is a Dynamic Antigen-Driven Differentiation Program Initiated Early during Tumorigenesis. Immunity 45, 389–401 (2016).

21. Anderson, K. G., Stromnes, I. M. & Greenberg, P. D. Cancer Cell Perspective Obstacles Posed by the Tumor Microenvironment to T cell Activity: A Case for Synergistic Therapies. Cancer Cell 31, 311–325 (2017).

22. Anz, D. et al. CD103 is a hallmark of tumor-infiltrating regulatory T cells. Int. J. Cancer 129, 2417–2426 (2011).

23. Ross, E. A. et al. CD31 is required on CD4 + T cells to promote T cell survival during Salmonella infection. J. Immunol. 187, 1553–1565 (2011).

24. Hanninen, A., Maksimow, M., Alam, C., Morgan, D. J. & Jalkanen, S. Ly6C supports preferential homing of central memory CD81 T cells into lymph nodes. Eur. J. Immunol. 41, 634–644 (2011).

25. Fourcade, J. et al. Upregulation of Tim-3 and PD-1 expression is associated with tumor antigen– specific CD8+ T cell dysfunction in melanoma patients. J. Exp. Med. 207, 2175–2186 (2010).

26. Mita, Y. et al. Crucial role of CD69 in anti-tumor immunity through regulating the exhaustion of tumor-infiltrating T cells. Int. Immunol. 30, 559–567 (2018).

27. Sun, C., Mezzadra, R. & Schumacher, T. N. Regulation and Function of the PD-L1 Checkpoint. Immunity 48, 434–452 (2018).

28. Bianchini, M. et al. PD-L1 expression on nonclassical monocytes reveals their origin and immunoregulatory function. Sci. Immunol. 4, eaar3054 (2019).

29. Busch, D. H., Pilip, I. M., Vijh, S. & Pamer, E. G. Coordinate regulation of complex T cell populations responding to bacterial infection. Immunity 8, 353–362 (1998).

30. Kaech, S. M. & Ahmed, R. Memory CD8 + T cell differentiation: initial antigen encounter triggers a developmental program in naïve cells. Nat. Immunol. 2, 415–422 (2001).

31. Jung, S. et al. In vivo depletion of CD11c+ dendritic cells abrogates priming of CD8+ T cells by exogenous cell-associated antigens. Immunity 17, 211–220 (2002).

32. Gabrilovich, D. I., Corak, J., Ciernik, I. F., Kavanaugh, D. & Carbone, D. P. Decreased antigen presentation by dendritic cells in patients with breast cancer. Clin. Cancer Res. 3, 483–490 (1997).

33. Schmid, P. et al. Atezolizumab and Nab-Paclitaxel in Advanced Triple-Negative Breast Cancer. N. Engl. J. Med. 379, 2108–2121 (2018).

34. Zuckerman, N. S. et al. Altered local and systemic immune profiles underlie lymph node metastasis in breast cancer patients. Int. J. Cancer 132, 2537–2547 (2012).

35. Wang, L. et al. Connecting blood and intratumoral Treg cell activity in predicting future relapse in breast cancer. Nat. Immunol. 20, 1220–1230 (2019).

36. Mittal, R., Wagener, M., Breed, E. R., Liang, Z. & Yoseph, B. P. Phenotypic T Cell Exhaustion in a Murine Model of Bacterial Infection in the Setting of Pre-Existing Malignancy. PLoS One 9, 93523 (2014).

37. Xie, J. et al. Pre-existing malignancy results in increased prevalence of distinct populations of CD4+ T cells during sepsis. PLoS One 13, e0191065 (2018).

38. Russ, A. J. et al. Melanoma-induced suppression of tumor antigen-specific T cell expansion is comparable to suppression of global T cell expansion. Cell. Immunol. 271, 104–109 (2011).

39. Klastersky, J. & Aoun, M. Opportunistic infections in patients with cancer. Ann. Oncol. 15, iv329–iv335 (2004).

40. Baluch, A. & Pasikhova, Y. Influenza Vaccination in Oncology Patients. Curr Infect Dis Rep 15, 486–490 (2013).

41. Danna, E. A. et al. Surgical Removal of Primary Tumor Reverses Tumor-Induced Immunosuppression Despite the Presence of Metastatic Disease. Cancer Res. 64, 2205–2211 (2004).

42. Kosaka, A., Ohkuri, T., Program, B. T. & Okada, H. Combination of an agonistic anti-CD40 monoclonal antibody and the COX-2 inhibitor celecoxib induces anti-glioma effects by promotion of type-1 immunity in myeloid cells and T-cells. Cancer Immunol Immunother 63, 847–857 (2014).

43. Tseng, W. W. et al. Cancer Therapy: Preclinical Development of an Orthotopic Model of Invasive Pancreatic Cancer in an Immunocompetent Murine Host. Clin. Cancer Res. 16, 3684–3695 (2010).

44. Kathryn E. Foulds, Lauren A. Zenewicz, Devon J. Shedlock, J. J. & Amy E. Troy, and H. S. Cutting Edge: CD4 and CD8 T Cells Are Intrinsically Different in Their Proliferative Responses. J Immunol 168, 1528–1532 (2002).

45. Spitzer, M. H. et al. An interactive reference framework for modeling a dynamic immune system. Immunology 349, 1259425.1–1259425.11 (2015).

46. Zunder, E. R. et al. Palladium-based Mass-Tag Cell Barcoding with a Doublet-Filtering Scheme and Single Cell Deconvolution Algorithm. Nat. Protoc. 10, 316–333 (2015).

47. Finck, R. et al. Normalization of mass cytometry data with bead standards. Cytom. Part A 83 A, 483–494 (2013).

48. Bair, E. & Tibshirani, R. Semi-Supervised Methods to Predict Patient Survival from Gene Expression Data. PLoS Biol. 2, 0511–0522 (2004).

49. Robert V. Bruggner, Bernd Bodenmiller, D. L. D. & Robert J. Tibshirani, and G. P. N. Automated identification of stratifying signatures in cellular subpopulations. PNAS 26, E2770–E2777 (2014).

50. Dumeaux, V. et al. Interactions between the tumor and the blood systemic response of breast cancer patients. PLoS Comput. Biol. 13, e1005680 (2017).

