## Supplementary Figures for Allen and Hiam et al. for "The Development, Function, and Plasticity of the Immune Macroenvironment in Cancer"

### Extended Data Fig. 1: Systemic immunity is distinctly remodeled across tumor types.

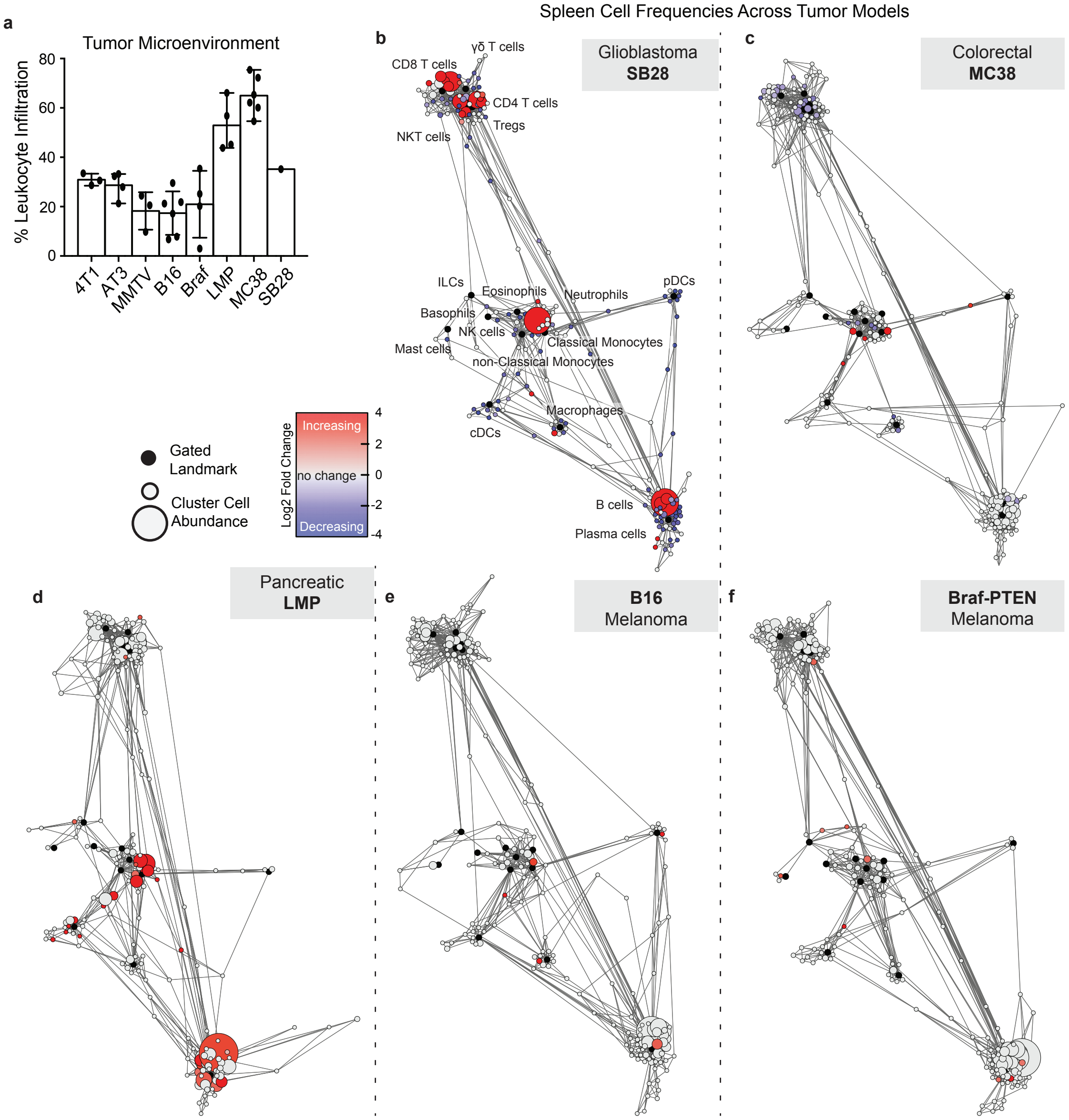

**Extended Fig. 2: Systemic immunity is distinctly remodeled over tumor development.**

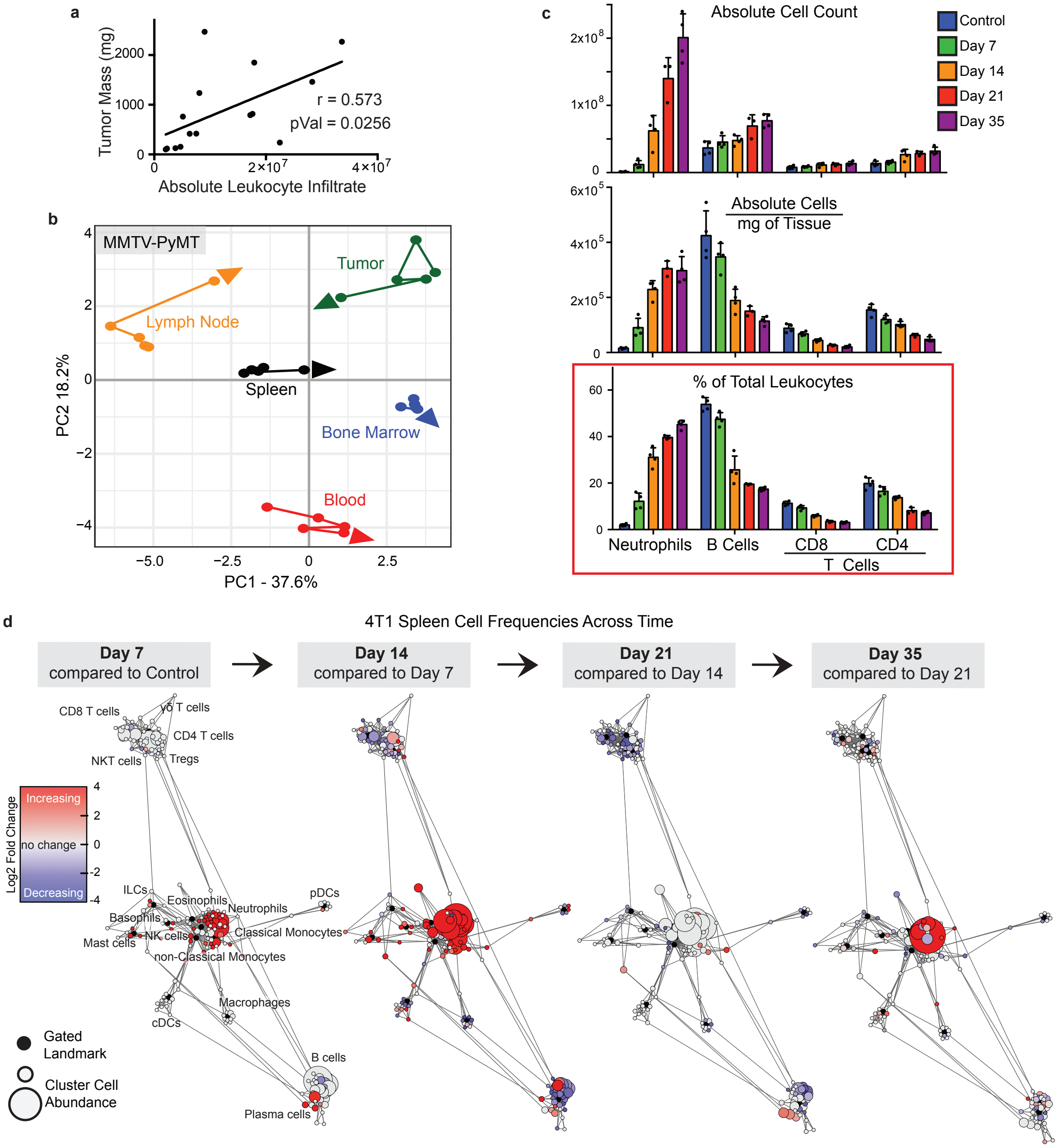

### Extended Fig. 3: Immunity is distinctly remodeled by compartment over tumor development.

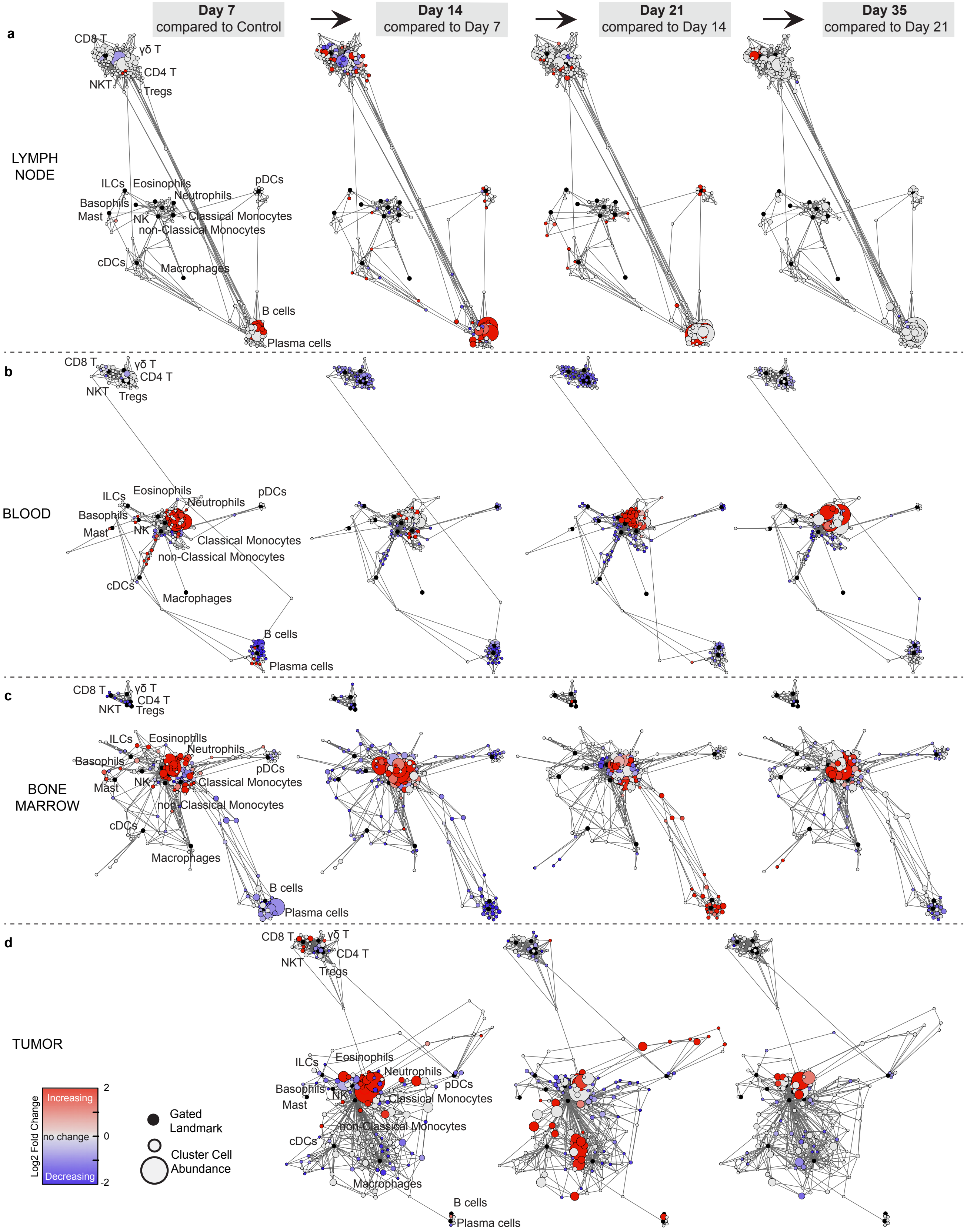

**Extended Fig. 4: Tumor growth shifts the systemic T cell composition across models.**

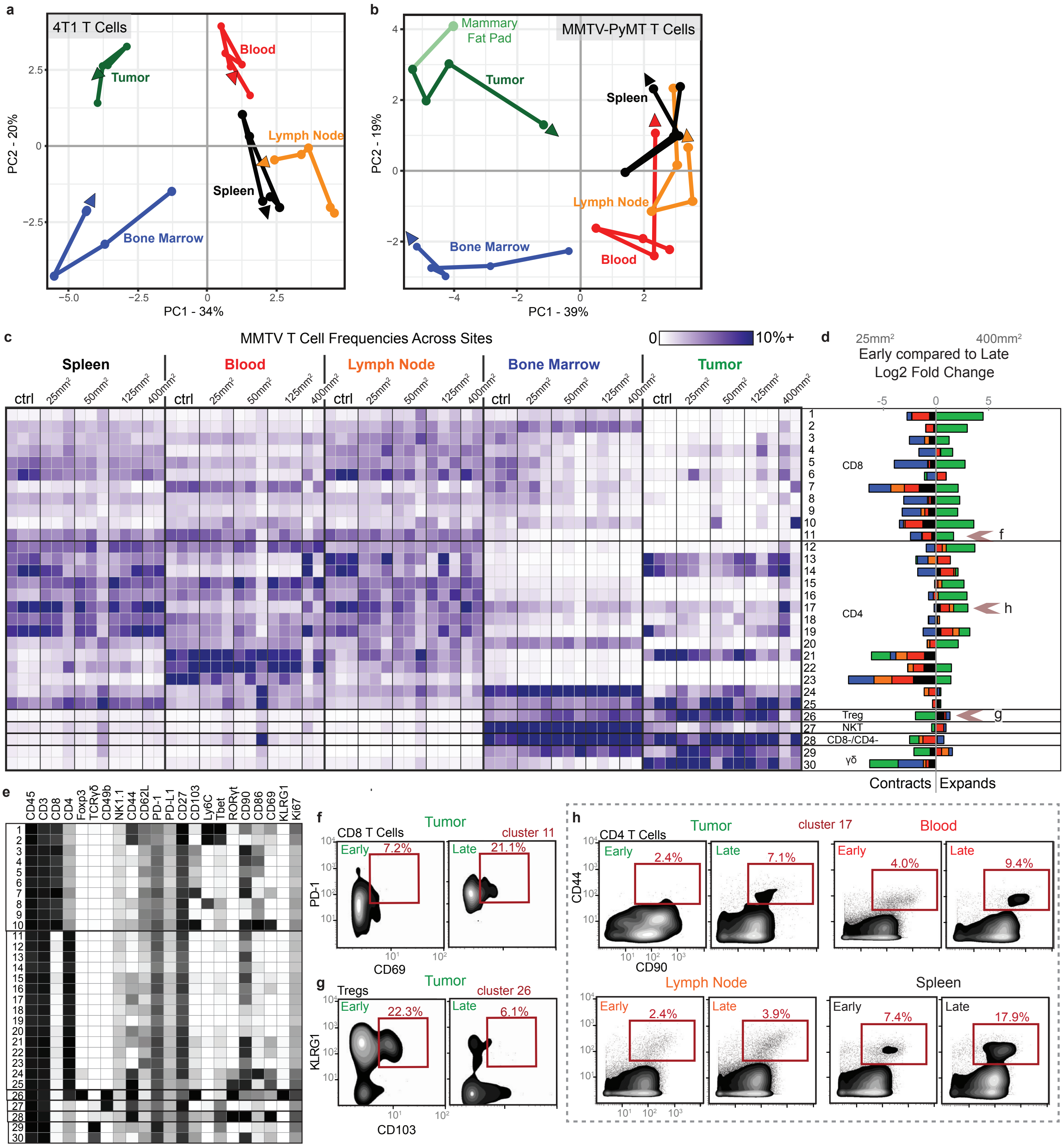

**Extended Data Fig. 5: Tumor growth shifts the systemic mononuclear phagocyte composition.**

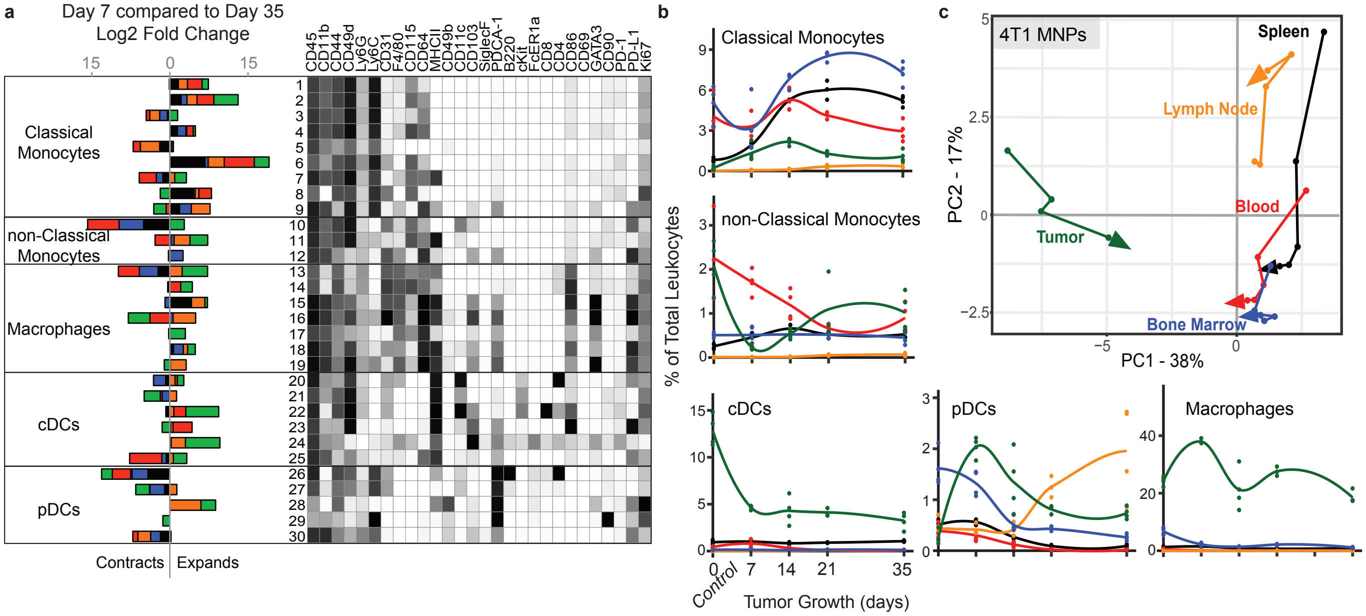

#### Extended Data Fig. 6: PD-1 and PD-L1 expression is dynamic over tumor growth.

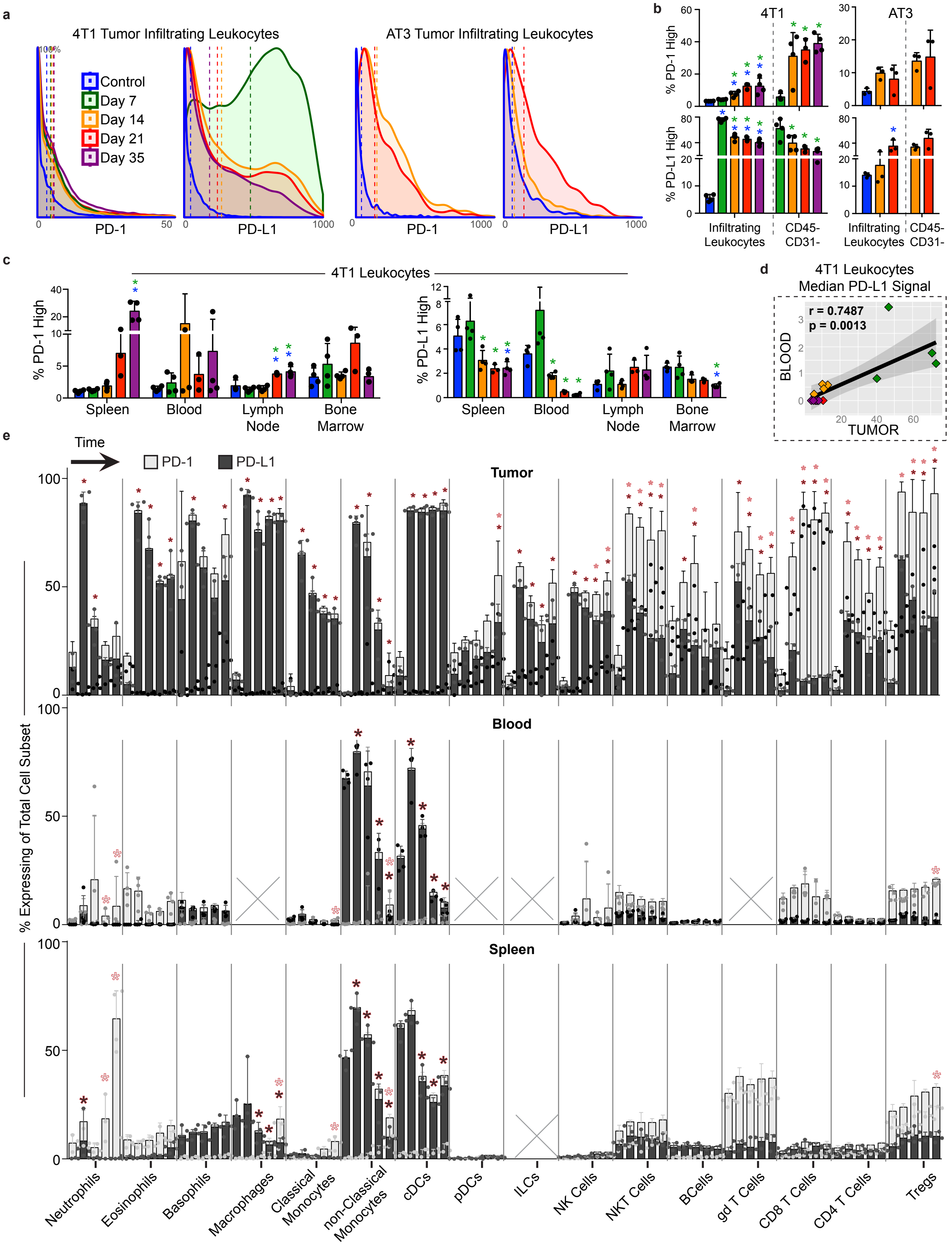

#### Extended Data Fig. 8: Tumor burden induces organ-specific changes in immune cell cycling.

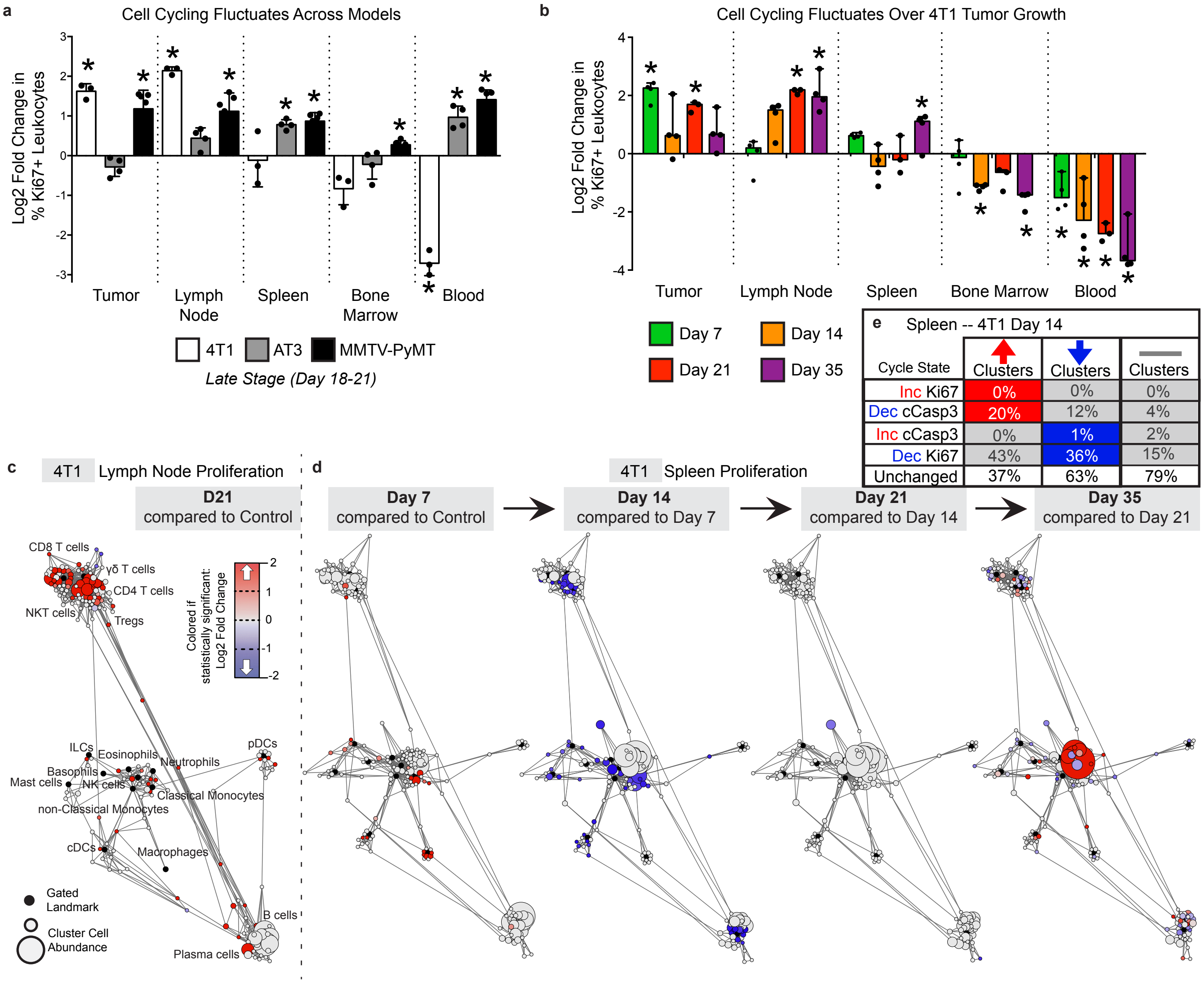


| Cycle State | ↑ Clusters | ↓ Clusters | — Clusters |
|---|---|---|---|
| Inc Ki67 | 0% | 0% | 0% |
| Dec cCasp3 | 20% | 12% | 4% |
| Inc cCasp3 | 0% | 1% | 2% |
| Dec Ki67 | 43% | 36% | 15% |
| Unchanged | 37% | 63% | 79% |

**Extended Fig. 8: Tumor driven deficits in T cell responses are cell-extrinsic.**

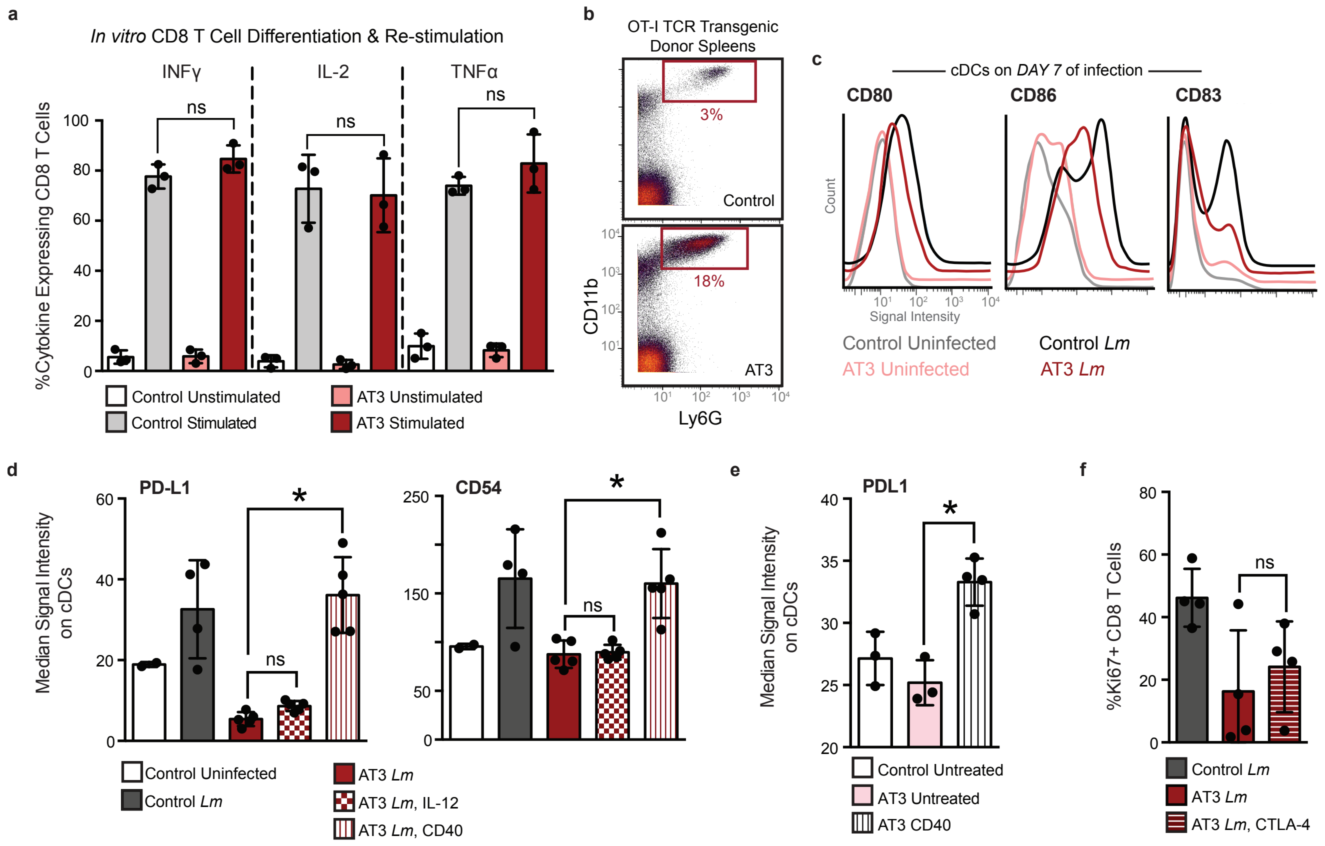

#### Extended Data Fig. 9: Tumor resection resets systemic immune organization and function.

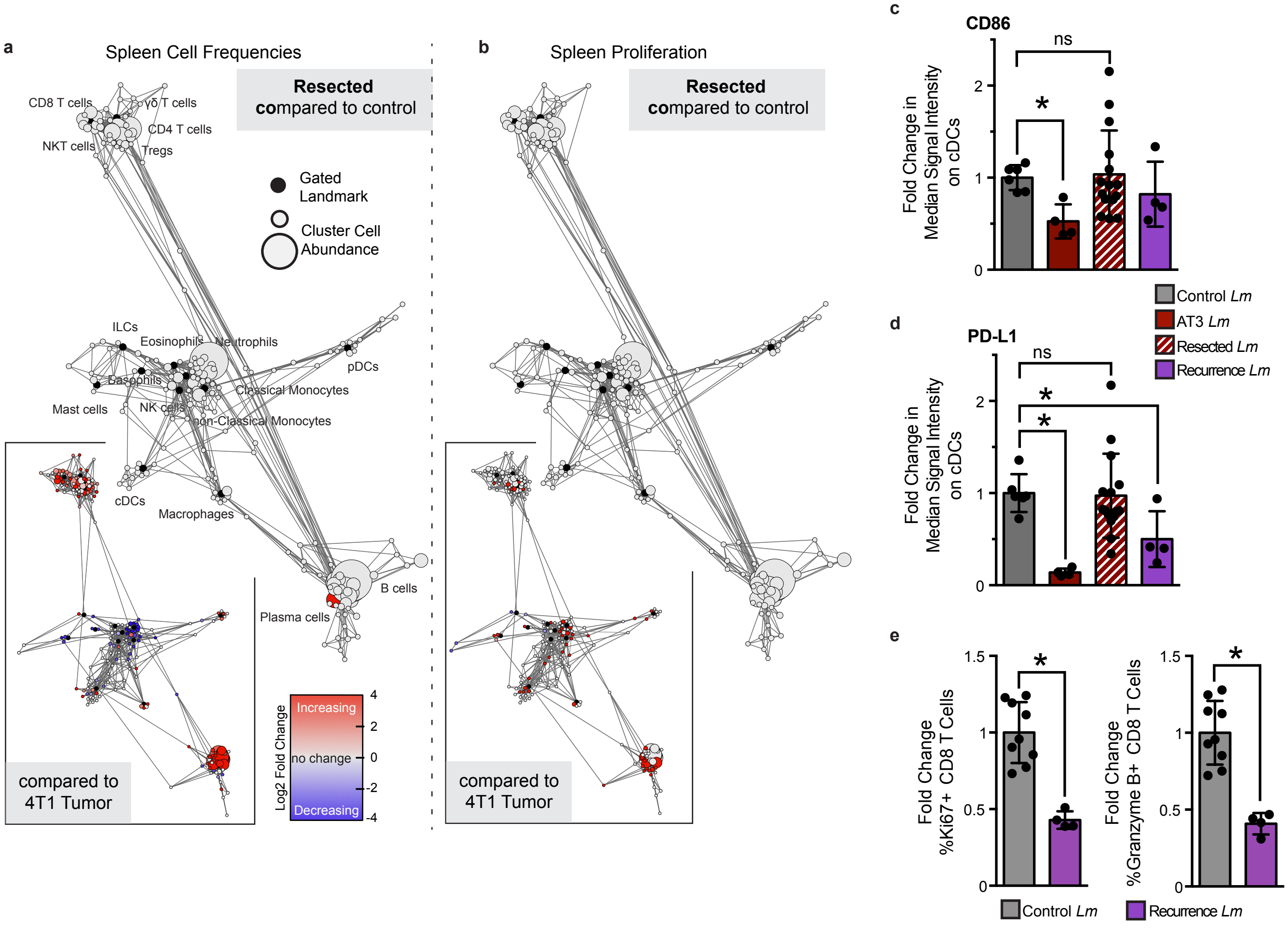
